# Accessory ESCRT-III proteins selectively regulate Rab11-exosome biogenesis in *Drosophila* secondary cells

**DOI:** 10.1101/2020.06.18.158725

**Authors:** Pauline P. Marie, Shih-Jung Fan, Claudia C. Mendes, S. Mark Wainwright, Adrian L. Harris, Deborah C. I. Goberdhan, Clive Wilson

**Affiliations:** Department of Physiology Anatomy and Genetics, University of Oxford; Department of Oncology, University of Oxford

## Abstract

Exosomes are secreted nanovesicles with potent signalling activity that are initially formed as intraluminal vesicles (ILVs) in multivesicular endosomes, which subsequently fuse with the plasma membrane. These ILVs are made in both late endosomes and recycling endosomes, the latter marked by the small GTPase Rab11 and generating exosomes with different cargos and functions. Core proteins within four Endosomal Sorting Complex Required for Transport (ESCRT) assemblies (0-III) play key sequential roles in late endosomal exosome biogenesis and ILV-mediated destruction of ubiquitinylated cargos through the endolysosomal system. They also control additional cellular processes, such as cytokinesis and other vesicle budding. By contrast, the functions of several accessory ESCRTs are not well defined. Here we assess the ESCRT-dependency of Rab11-exosomes, using RNA knockdown in *Drosophila* secondary cells (SCs) of the male accessory gland, which have unusually enlarged Rab11-positive compartments. Unexpectedly, not only are core proteins in all four ESCRT complexes required for Rab11-exosome formation, but also accessory ESCRT-III proteins, CHMP1, CHMP5 and IST1. Suppressing expression of these accessory proteins does not affect other aspects of cell morphology, unlike most core ESCRT knockdowns, and does not lead to accumulation of ubiquitinylated cargos. We conclude that accessory ESCRT-III components have a specific and potentially ubiquitin-independent role in Rab11-exosome generation, which might provide a target for blocking the pro-tumorigenic activities of these vesicles in cancer.

## Introduction

Exosomes are small extracellular vesicles (EVs) initially generated as intraluminal vesicles (ILVs) in endosomal multivesicular endosomes (MVEs) (Colombo et al., 2014; Möbius et al., 2002; White et al., 2006). They are released by all cell types and mediate cell-cell communication events in development, immunity, reproduction and many other physiological and disease processes, such as neurodegenerative diseases (Thompson et al., 2016) and cancer (Saber et al., 2020). In cancer, exosome cargos can modify the behaviour of recipient cells, leading to tumour progression by driving tumour cell growth and invasiveness, reprogramming the tumour microenvironment, promoting endothelial network assembly, modulating the immune response and inducing pre-metastatic niche formation (Becker et al., 2016; Comito et al., 2020; Saber et al., 2020). Exosomes are highly heterogeneous, but the mechanisms underlying this heterogeneity and the routes by which exosome signalling can be modulated have until recently, remained largely unexplored.

MVEs are typically assumed to be of late endosomal origin. At least two exosome biogenesis mechanisms have been reported in these compartments. One involves the Endosomal Sorting Complex Required for Transport (ESCRT) proteins, a modular ILV-generation system originally discovered in yeast (Raymond et al., 1992). The other ESCRT-independent mechanism requires ceramide (Trajkovic et al., 2008). The ESCRT machinery is composed of four protein subcomplexes named ESCRT-0, -I, -II, and –III, and the ATPase Vps4. These assemble sequentially at the limiting membrane of the late endosome (Schuh and Audhya, 2014; Vietri et al., 2019). The ESCRT machinery regulates various cellular processes involved in development (Irion and St Johnston, 2007), neurogenesis (Loncle et al., 2015), virus budding (Carlton et al., 2008), cytokinesis (König et al., 2017; Matias et al., 2015) and autophagy (Lee et al., 2007; Rusten et al., 2007). ESCRT malfunction is also associated with numerous human diseases (for review see Saksena and Emr 2009). Recently, two distinct cancer exosome populations with differing cargos have been reported following asymmetric-flow fractionation. Although they share some classic exosomes markers, small (S) exosomes are enriched in tetraspanins, whereas large (L) exosomes selectively accumulate G-proteins and integrins, suggesting different functions (Jimenez et al., 2018). However, the cellular mechanisms generating these two exosome subtypes remain uncharacterised.

ESCRT-III proteins are subdivided into core and accessory molecules. The function of the latter in metazoans is not well understood. *Chmp1, Ist1, Vta1* and *Chmp5* appear to be required for the full function of Vps4 in yeast endosomes. Chmp1 associates with Ist1, and Vta1 with Chmp5 in two distinct sub-complexes (Nickerson et al., 2010). Single accessory ESCRT-III knockdowns or mutants often have no or very weak phenotypes (Agromayor et al., 2009; Dimaano et al., 2008; Rue et al., 2007), whereas *Vps4* nulls produce severe phenotypes (Babst et al., 1998), impairing ESCRT complex dissociation. Some combinations of accessory ESCRT-III mutations lead to formation of vesicle-free endosomes in yeast (Nickerson et al., 2010) and *Drosophila* (Bäumers et al., 2019), consistent with loss of Vps4 function. This suggests that accessory ESCRT-III proteins may often be individually dispensable, but they provide robustness to the ESCRT pathway under stress conditions, for example, when cells are highly metabolically active (Shim et al., 2006; Agromayor et al., 2009), or at low temperatures (Bäumers et al., 2019).

We have recently shown in both human cancer cell lines and in prostate-like secondary cells (SCs) of the male accessory gland in the fruit fly, *Drosophila melanogaster*, that exosomes are not only generated in late endosomes, but also in recycling endosomes marked by the trafficking regulator Rab11 (Fan et al., 2019). These latter exosomes, some of which are marked by Rab11, have different cargos and functions to late endosomal exosomes, and in cancer cells, their secretion is induced by metabolic stresses via suppression of nutrient-sensitive mechanistic Target of Rapamycin Complex 1 (mTORC1) signalling.

In this study, we use *Drosophila* SCs to perform an extensive genetic screen of the ESCRT proteins for roles in Rab11-exosome biogenesis and secretion. We show that all four ESCRT sub-complexes are involved in these processes. Importantly, we establish that accessory ESCRT-III genes are also essential for the normal biogenesis of Rab-11 exosomes, but unlike other ESCRTs, are not required for the ILV-dependent mechanism that removes ubiquitinylated proteins from the surface of late endosomes. Our data therefore demonstrate a unique role for accessory ESCRT-III proteins in Rab11-exosome biogenesis, which does not appear to require the ubiquitinylation-dependent cargo loading associated with endolysosomal trafficking.

## Results

### ESCRT sub-complexes 0-II are involved in maturation of large non-acidic compartments and ILV biogenesis within them

The *Drosophila* accessory gland is composed of two cell types, the main cells and the secondary cells (SCs; Fig 1A). SCs have unusually large secretory and endosomal compartments, including between three and five large (> 2 µm) Rab7-positive late endosomes and lysosomes (LELs), which can be visualised with LysoTracker® and depending on the marker used, between ten and twenty large (> 2 µm) non-acidic compartments (Corrigan et al., 2014; Fan et al., 2019) Fig 1F,G). The latter can be fluorescently marked by a GFP-tagged form of Breathless (Btl), a fly FGF receptor homologue, when overexpressed in SCs, or by a *YFP-Rab11* gene trap (Fig 1B-E). Inside these compartments, Btl-GFP and YFP-Rab11 also label ILVs (Fig 1D’, E’ Zoom) (Fan et al., 2019), collectively called Rab11-exosomes, with some particularly bright puncta often observed for Btl-GFP. These puncta typically surround a central, protein-rich dense-core granule (DCG), which can be visualised with differential interference contrast microscopy (DIC; Fig 1B-E; see top panels in Fig 1D’,E’). The markers also label secreted exosomes in the AG lumen (Fig 1D’’ (Fan et al., 2019), when visualised as shown in Fig 1A’’.

**Figure 1.**
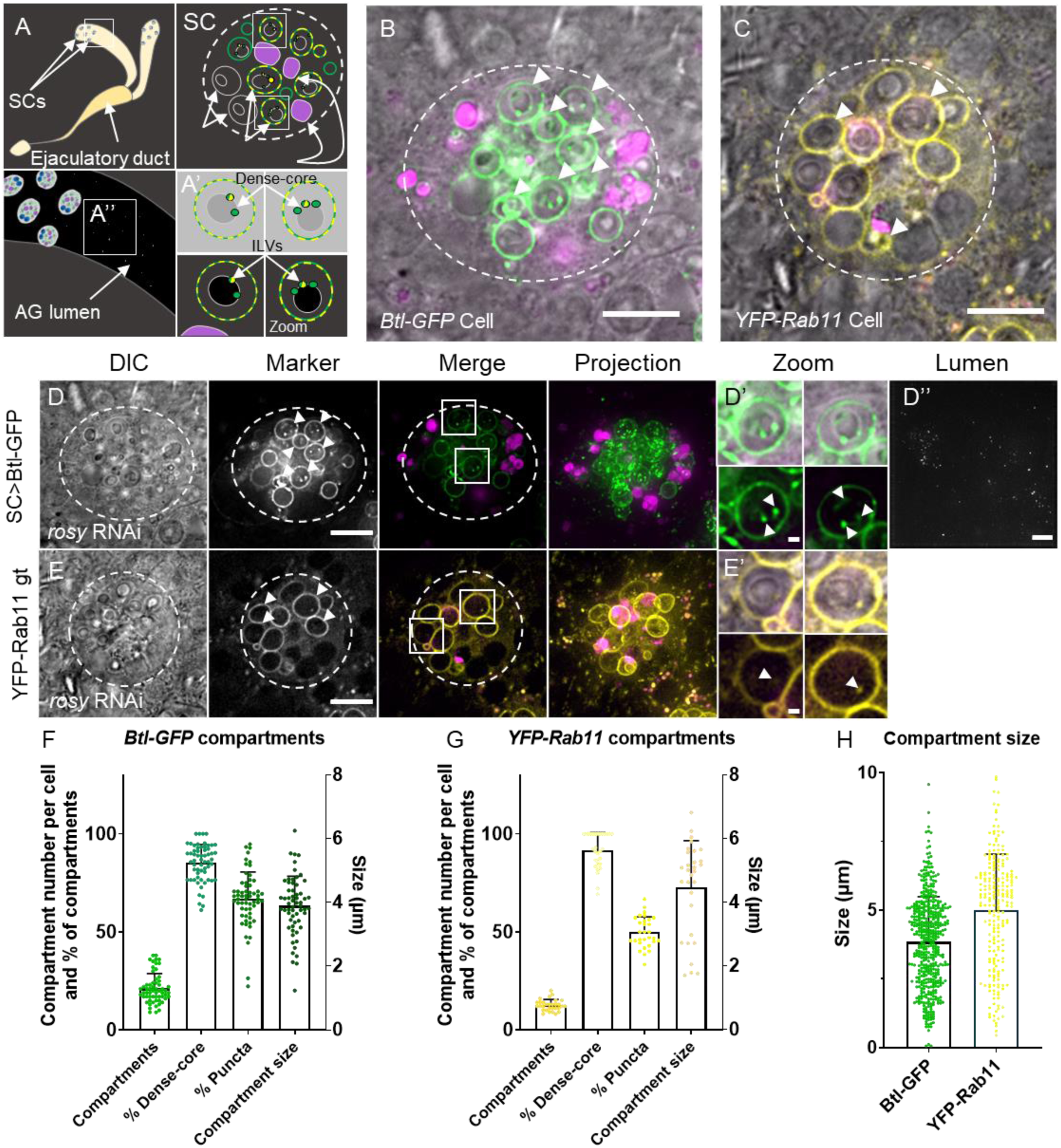
Large non-acidic compartments of *Drosophila* SCs form DCGs and exosomes. (A) Schematic of accessory gland (AG), marking position of secondary cells (SCs) at distal tip (top left panel) and schematic representation of a basal Z-plane view of a single SC (top right), which corresponds to SC images in all figures. SC schematic highlights large non-acidic compartments in green and yellow, and late endosomes and lysosomes in magenta. The outline of SCs is approximated by a dashed circle. As in the images in D-E, boxed non-acidic compartments are magnified in two zooms (with DIC to visualise DCG [top], and without [bottom]; A’), to show ILVs, a subset of which are marked by Rab11. Finally, the central region of the lumen imaged in the Lumen column (D’’) is shown schematically (A’’; 10 µm projection with 0.3 µm spacing). (B) SC expressing the Btl-GFP marker. (C) Cell expressing the *YFP-Rab11* gene trap. (D) SC expressing control *rosy*-RNAi and Btl-GFP marker. Btl-GFP puncta are observed inside large non-acidic compartments (arrowheads; D’ Zoom), which contain DCGs (visible by DIC), and in the AG lumen (D’’). (E) SC from *YFP-Rab11* gene trap male expressing *rosy*-RNAi. YFP-Rab11 puncta are observed inside large Rab11-positive compartments (arrowheads; E’ Zoom), which contain DCGs (visible by DIC). (F) Bar chart showing number of Btl-GFP-positive compartments per cell, percentage of them containing DCGs and Btl-GFP-positive ILVs, and average size of compartments per cell. n = 30 SCs. (G) Bar chart showing number of YFP-Rab11 compartments per cell, percentage of them containing Dense-core granule, YFP-Rab11-positive ILVs and average size of compartments per cell. n = 30 SCs. (H) Individual non-acidic compartment sizes in SCs expressing the two markers. n = 30 SCs. All data are from 6-day-old males shifted to 29°C at eclosion to induce expression of transgenes. Genotypes are: *w; P[w*^*+*^, *tub-GAL80ts]/+; dsx-GAL4/P[w*^*+*^, *UAS-btl-GFP]* or *w; P[w*^*+*^, *tub-GAL80ts]/+; dsx-GAL4/ w; TI{TI}Rab11EYFP* with or without UAS-rosy-RNAi expression. Scale bars in B-D and Lumen column, 10 µm; in D’, E’ Zoom, 1 µm.

In previous work, we found that Btl-GFP and YFP-Rab11 mark approximately the same number of large non-acidic compartments with diameter > 2 µm (Fan et al., 2019). However, we observed that a stable fly line permitting inducible overexpression of Btl-GFP also marked multiple smaller (0.4-2 µm diameter; Fig 1H) non-acidic compartments, so that the total number of labelled non-acidic secretory compartments was 21 ± 8, compared to 13 ± 3 with YFP-Rab11 (Fig 1F). 67 ± 14 % of these compartments contained Btl-GFP-positive ILVs and 85 ± 9% contained DCGs (Fig 1B, D, F), compared to 50 ± 12 % and 87 ± 11 % respectively in a *YFP-Rab11* background; Fig 1C, E, G) demonstrating that most non-acidic compartments are DCG- and exosome-containing secretory compartments.

We previously showed that ILV formation inside large non-acidic SC compartments requires the ESCRT-0 Stam, ESCRT-I Vps28 and ESCRT-III Shrub (Fan et al., 2019). The roles of other ESCRT sub-complex components in Rab11-exosome formation and secretion, including the accessory ESCRT-III proteins, were tested.

Flies were generated that expressed RNAis targeting genes encoding members of all four ESCRT classes in adult SCs, using the GAL4/UAS system under temperature-sensitive GAL80^ts^-inducible control (Fan et al., 2019). At least two genes in each ESCRT family were knocked down, typically by expressing in different crosses two independent RNAis, immediately after adult eclosion.

The first step of exosome biogenesis involves the ESCRT-0 proteins, Hrs and Stam (respectively Vps27 and Hse1 in yeast). As in mammalian cells (Razi and Futter, 2006), SCs depleted of ESCRT-0 components, which cluster mono-ubiquitinylated cargos at sub-domains on the endosome membrane (Prag et al., 2007), had enlarged acidic compartments (Fig 2D, S1B), when compared to SCs expressing a control RNAi targeting *rosy* (Fig 2C, S1A). *rosy* encodes the enzyme xanthine dehydrogenase required for normal eye colour, which has no known role in SC secretion. For some RNAi lines, *ESCRT-0* knockdown cells contained more than twice as many Btl-GFP-positive compartments with diameter > 0.4 µm, when compared to controls (Fig 2A, 2C, 2D, S1B, S2A). However, a much-reduced proportion of these compartments formed ILVs (Fig 2B, 2C’, 2D’, S1B’, S2B). Most *Hrs* knockdown SCs lacked DCG-containing compartments, although this phenotype was not observed with *Stam*-RNAi (Fig 2H). Furthermore, secretion of Btl-GFP puncta was also greatly decreased (Fig 2D’’, S1B’’, 2I, S2C).

**Figure 2.**
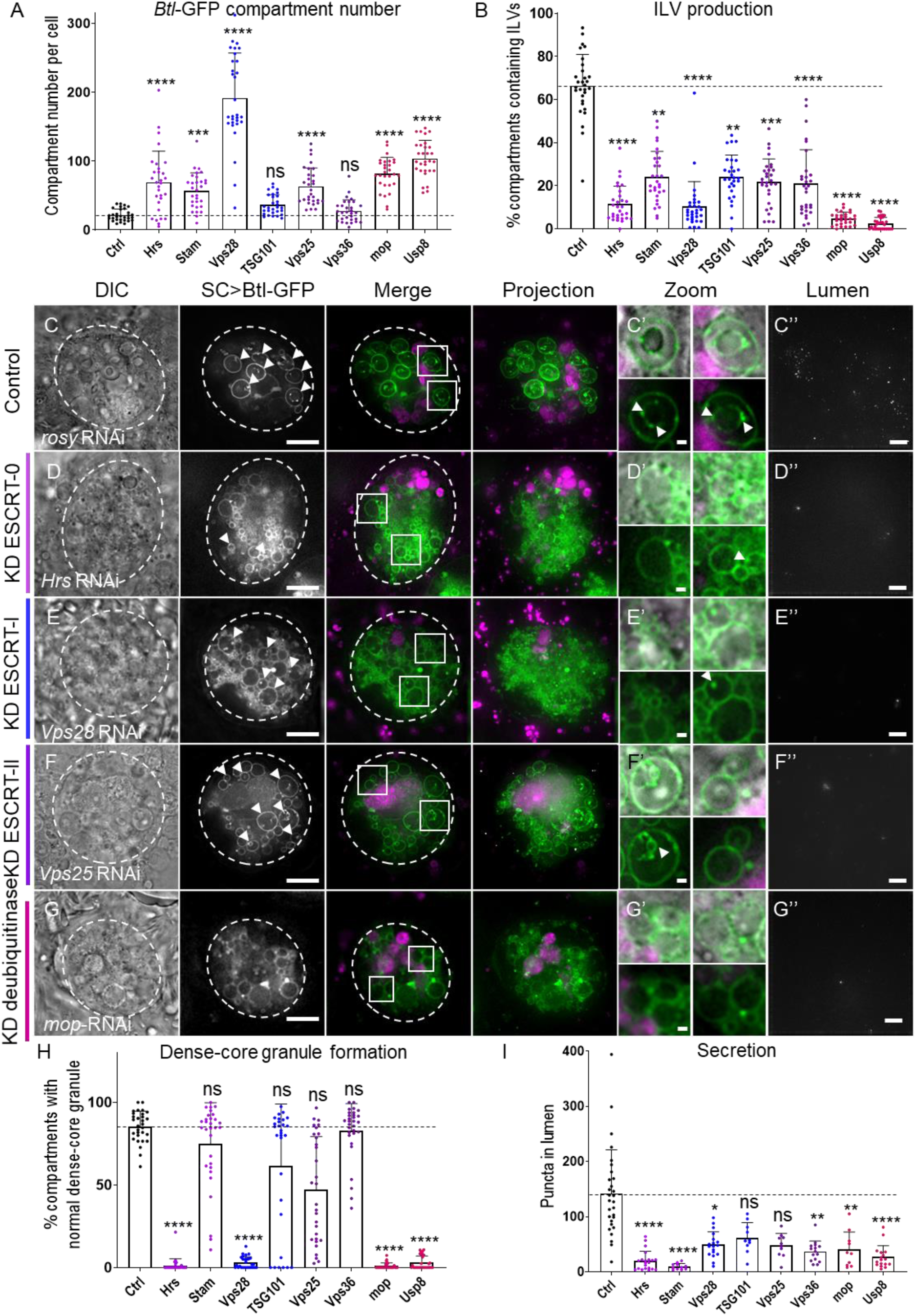
ESCRT-0-II components regulate exosome biogenesis in non-acidic compartments of *Drosophila* SCs. (A) Bar chart showing the number of non-acidic Btl-GFP-positive compartments per SC with diameter > 0.4 µm in control versus ESCRT-0, -I, -II and deubiquitinylases *mop* and *Usp8* knockdowns (n = 30). (B) Bar chart showing percentage of Btl-GFP compartments containing Btl-GFP-positive ILVs. n = 30 SCs. Panels C-G show basal wide-field fluorescence views of living SCs from 6-day-old males expressing a *UAS-Btl-GFP* (green) and a *UAS-RNAi* construct under temperature-induced, SC-specific GAL4 control from eclosion onwards. SC outline is approximated by dashed white circles. Acidic compartments are marked by LysoTracker® Red (magenta). Boxed non-acidic compartments are magnified in C’-G’ Zooms. Transverse images of AG lumens are shown in C’’-G’’. (C) SC expressing Btl-GFP and *rosy-*RNAi construct (control). Btl-GFP-positive ILVs are observed inside large non-acidic compartments (arrowheads; C’) and as puncta in AG lumen (C’’). DCGs are visible in mature Btl-GFP-positive compartments (DIC and C’). (D) SCs expressing *Hrs-*RNAi lack DCGs (DIC) and have a reduced proportion of Btl-GFP-positive ILV-containing compartments (D’) and secreted puncta (D’’). (E) SCs expressing *Vps28*-RNAi also lack DCGs and have reduced exosome biogenesis. (F) SCs expressing *Vps25*-RNAi form DCGs and enlarged ILVs in some non-acidic compartments (F’). Exosome secretion is reduced. (G) SCs expressing *mop*-RNAi lack DCGs (DIC) and have a reduced proportion of Btl-GFP-positive ILV-containing compartments. Exosome secretion is reduced. (H) Bar chart showing proportion of Btl-GFP-positive compartments containing DCGs. n = 30 SCs. (I) Bar chart showing number of Btl-GFP fluorescent puncta in the lumen of AGs with ESCRT knockdown compared to control. n > 10. All data are from 6-day-old males shifted to 29°C at eclosion to induce expression of transgenes. Genotypes are: w; *P[w*^*+*^, *tub-GAL80ts]/+; dsx-GAL4/P[w*^*+*^, *UAS-btl-GFP]* with *UAS-rosy-RNAi* knockdown construct (C), *UAS-Hrs-RNAi*-#1 (D), *UAS-Stam-RNAi*-#1, *UAS-Vps28-RNAi*-#1 (E), *UAS-TSG101-RNAi*-#1, *UAS-Vps25-RNAi*-#1 (F), *UAS-Vps36-RNAi*-#1, *UAS-mop-RNAi*-#1 (G), *UAS-Usp8-RNAi*-#1. Scale bars in C-G and C’’-G’’, 10 µm; in C’-G’, 1 µm. Data were analysed by Kruskal-Wallis test. *p < 0.05, **p < 0.01, ***p < 0.001 and ****p < 0.0001 relative to control. See also Fig S1 for images of gene knockdowns not depicted here and Fig S2 for additional RNAi lines tested (#2).

ESCRT-0 proteins recruit the ESCRT-I and ESCRT-II complexes, which deform the membrane into cargo-containing vesicles (Henne et al., 2011; Huotari and Helenius, 2011). Knockdown of the ESCRT-I *Vps28* produced a striking increase in the number of Btl-GFP-positive compartments (Fig 2A, S2A), which were almost all devoid of internal puncta (Fig 2B, 2E, S2B). Knockdown of a second ESCRT-I component *TSG101* (*Vps32* in yeast), an established marker of late endosomal exosomes (Kumar et al., 2014), also reduced the proportion of ILV-containing Btl-GFP-marked compartments and like *Vps28*-RNAi, decreased secretion (Fig 2B, 2I, S1C, S2A-C). However, it did not significantly reduce the number of SCs containing DCGs, in sharp contrast to *Vps28* knockdown (Fig 2H).

Compared to other ESCRTs, knockdown of ESCRT-II components *Vps25* and *Vps36* had a milder effect on Btl-GFP-labelled compartments (Fig 2F, S1D). Compartment size was comparable to controls and for both *Vps36* knockdowns, compartment number was not significantly increased (Fig 2A, S2A). The proportion of ILV-producing compartments decreased significantly with only one of the two knockdowns for each *ESCRT-II* gene (Fig 2B, S2B). ILVs, when present, were often enlarged (Fig 2F’, S1D’). DCG formation was not affected (Fig 2H), and although exosome secretion generally appeared reduced, the effects were not significant (Fig 2I and Fig S2C).

Usp8 and Mop are deubiquitinylases known to be involved in ESCRT-dependent, late endosomal ILV production. Usp8, which deubiquitinates and hence stabilizes Hrs, is also involved in the deubiquitinylation of EV cargo before internalization (Ali et al., 2013; Alwan and Leeuwen, 2007), while Mop inhibits the ubiquitin ligase Cbl, which represses Hrs, and therefore blocks the destabilisation of Hrs (Pradhan-Sundd and Verheyen, 2015). *Usp8* and *mop* knockdown both produced phenotypes similar to *ESCRT-0* knockdowns (Fig 2G, S1E), with increased numbers of small Btl-GFP-positive compartments, most of which did not produce ILVs (Fig 2A, 2B, S2A, S2B). As with *Hrs* knockdown, most cells did not produce dense cores inside their secretory compartments (Fig 1H). A strong reduction in secretion was observed for these two deubiquitinases (Fig 2I, S2C).

Overall, we conclude that ESCRT-0, -I and -II components all regulate the biogenesis of exosomes in large non-acidic SC compartments, as well as in late endosomes, though the effects of *ESCRT-II* knockdowns seem milder than other ESCRTs. Interestingly, there is also a variable effect on DCG formation, even within the same ESCRT subclass, although generally knockdowns with the strongest effects on ILV formation are those that suppress DCG biogenesis.

### Both core and accessory ESCRT-III proteins are involved in Rab11-exosome biogenesis

We investigated the roles of the core ESCRT-III proteins, which coordinate the membrane-severing function in late endosomal ILV formation. First, Vps20 mediates the recruitment and oligomerization of Shrub (Snf7 in yeast, CHMP4a-c in mammals), the major component of the ESCRT-III complex (Babst et al., 2002a, 2002b; Howard et al., 2001), which forms a helical filament that triggers membrane constriction. Vps24 caps the ESCRT-III filament and Chmp2 (Vps2 in yeast) recruits the AAA ATPase Vps4 to promote scission of the vesicle neck (Henne et al., 2012; Teis et al., 2010).

Consistent with previous work (Fan et al., 2019), knockdown of *shrub* produced a strong phenotype; cells were enlarged and full of very small compartments lacking ILVs (Fig 3B, G-H, S2D-E). LELs were either extremely small or absent, and DCGs were not detectable (Fig 3I). Secretion was also strongly impaired (Fig 3J, S2F). Knockdown of the core ESCRT-III genes *Chmp2* (Fig 3C) and*Vps20* (Fig S1F) produced similar, but weaker, effects (Fig 3G-J, S2D-F). Knockdown of *Vps24* had even milder effects inside SCs, although it did significantly reduce exosome secretion and have a partial effect on DCG formation (Fig 3G-J, S1G, and S2D-F).

**Figure 3.**
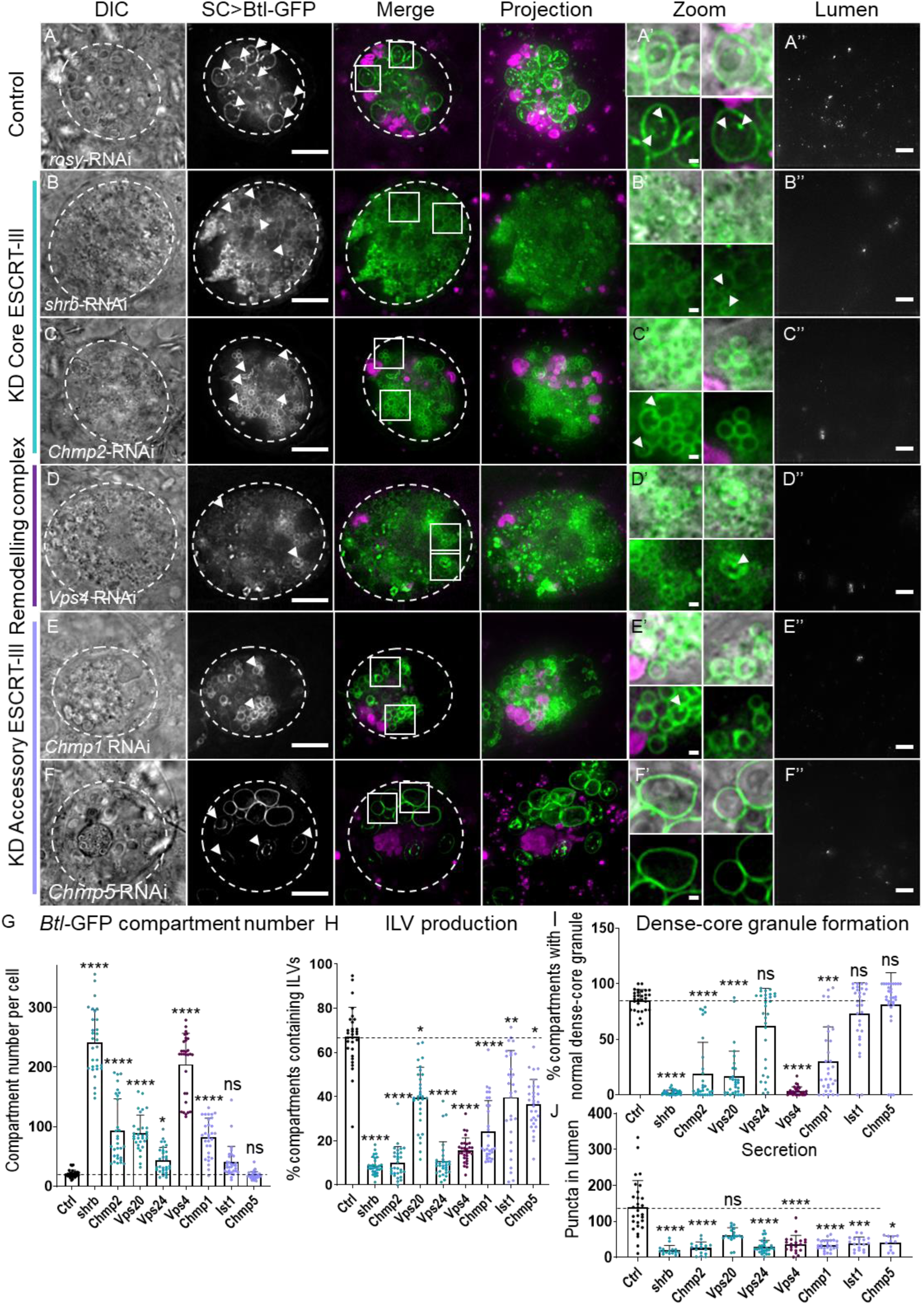
Both core and accessory ESCRT-III components regulate exosome biogenesis in non-acidic compartments of SCs. Panels A-F show basal wide-field fluorescence views of living SCs expressing Btl-GFP (green). SC outline approximated by dashed white circles. Acidic compartments are marked by LysoTracker® Red (magenta). Boxed non-acidic compartments are magnified in A’-F’ (Zoom). Transverse image projections of AG lumens are shown in A’’-F’’. (A) Control SC expressing *rosy*-RNAi. Btl-GFP-positive vesicle membranes are observed inside large non-acidic compartments (arrowheads; A’) and as puncta in AG lumen (A’’). (B) SC expressing *shrub-*RNAi. Number of non-acidic compartments is greatly increased. No large LELs or DCGs are observed. Btl-GFP-positive ILVs and puncta in the AG lumen are strongly reduced. (C) SC expressing *Chmp2-*RNAi. Non-acidic compartment number is increased. There are few DCGs and Btl-GFP-positive ILVs and puncta in the AG lumen are reduced. (D) SC expressing *Vps4*-RNAi. Btl-GFP-positive compartments are greatly increased. Btl-GFP-positive ILVs and puncta in the AG lumen are reduced. (E) SC expressing *Chmp1-*RNAi. Non-acidic compartment number is increased. Btl-GFP-positive ILVs and puncta in the AG lumen are reduced. (F) SC expressing *Chmp5*-RNAi. Btl-GFP-positive ILVs and puncta in the AG lumen are reduced. (G) Bar chart showing the number of non-acidic Btl-GFP-positive compartments with diameter > 0.4 µm per SC in control vs *ESCRT-III* knockdowns. n = 30. (H) Bar chart showing percentage of Btl-GFP-compartments containing Btl-GFP-positive ILVs. n = 30 SCs. (I) Bar chart showing proportion of Btl-GFP-positive compartments containing DCGs, visible by phase contrast imaging. n = 30 SCs. (J) Bar chart showing Btl-GFP fluorescent puncta number in the lumen of AGs with ESCRT-III knockdowns compared to controls. n > 10 AG lumens. All data are from 6-day-old male flies shifted to 29°C at eclosion to induce expression of transgenes. Genotypes are: w; *P[w+, tub-GAL80ts]/+; dsx-GAL4/P[w+, UAS-btl-GFP]* with *UAS-rosy-RNAi* knockdown construct (A), *UAS-shrb-RNAi*-#1 (B), *UAS-Chmp2*-*RNAi*-#1 (C), *UAS-Vps20*-*RNAi*-#1, *UAS-Vps24-RNAi*-#1, *UAS-Vps4*-*RNAi*-#1 (D) *UAS-Chmp1-RNAi*-#1 (E), *UAS-Ist1*-*RNAi*-#1, *UAS-Chmp5-RNAi*-#1 (F). Scale bars in A-F and A’’-F’’, 10 µm, and in A’-F’, 1 µm. Data were analysed by Kruskal-Wallis test. *p < 0.05, **p < 0.01, ***p < 0.001 and ****p < 0.0001 relative to control. See Fig S1 for images of gene knockdowns not depicted here, and Fig S2 for additional RNAi lines tested.

*Vps4* knockdown produced similar effects to blocking the principal ESCRT-III proteins. Hundreds of small compartments, most without internal puncta, were seen, and only small LELs were present, while secretion of Btl-GFP was strongly suppressed, and dense cores were absent (Fig 3D, G-J, S2D-F).

Accessory ESCRT-III functions remain elusive. They have been described as acceptors, modulators and enhancers for ESCRT-III-mediated vesicle scission and Vps4 dissociation (Rue et al., 2007; Azmi et al., 2008). Very few studies have revealed phenotypes with single accessory ESCRT-III mutations or knockdowns (Shim et al., 2006; Loncle et al., 2015; Frankel et al., 2017).

Surprisingly, knockdown of the other accessory *ESCRT-III* genes also significantly affected ILV formation in Btl-GFP-positive SC compartments. C*hmp1* knockdown produced a phenotype comparable to *Chmp2*-RNAi (Fig 3E, G-J and S2D-F). Knockdown of *Chmp5* and *Ist1*, two other accessory ESCRT-III components had milder effects on compartment size and number, and DCGs, but did reduce the proportion of compartments with ILVs and exosome secretion significantly (Fig 3E-J, Fig S1H, S2D-F).

In summary, core and accessory ESCRT-III proteins are involved in exosome biogenesis in non-acidic compartments of SCs (Fig 3H, J, S2E-F). Knockdown of several core ESCRT-III genes, like *shrb* and *Vps4*, increases the number and reduces the size of large non-acidic compartments marked with Btl-GFP, which frequently lack ILVs and a DCG. By contrast, reducing expression of the accessory ESCRT-III proteins, Chmp5 and Ist1, does not have major effects on SC compartment organisation, but still affects ILV formation and exosome secretion.

### *ESCRT* knockdowns do not affect non-acidic compartment identity in *Drosophila* secondary cells

One possible explanation for the reduction of ILV-containing DCG compartments and associated defective DCG formation following *ESCRT* knockdown is that the identity of these compartments is altered by these genetic manipulations. To investigate this, a YFP-tagged ‘gene trap’ marking Rab11 at the endogenous *Rab11* locus in *Drosophila* (Dunst et al., 2015) was used to determine whether the defective compartments carry Rab11 at their limiting membrane and whether Rab11 marks a subset of ILVs, as is observed in wild-type cells (Fan et al., 2019). SCs in this gene trap line have essentially normal compartmental organisation (12 ± 3 Rab11-positive compartments per cell; Fig 1G) and their morphology is generally less disturbed by *ESCRT* knockdown than in SCs expressing other ILV markers (Fan et al., 2019).

Control SCs contain 10-15 large DCG-containing, non-acidic, Rab11-positive secretory compartments (Fig 1E,G, Fig 4A-C) and 3-4 LysoTracker®-positive LELs (Fig 4H). About 50% of Rab11-compartments contain fluorescent ILVs around the DCG (Fig 4B), including scattered bright puncta, representing a subset of ILVs present (Fig 4C; arrowheads in Fig 4C’). Although these ILVs are secreted into the lumen of the gland, the low levels preclude quantification of secretion with this marker.

**Figure 4.**
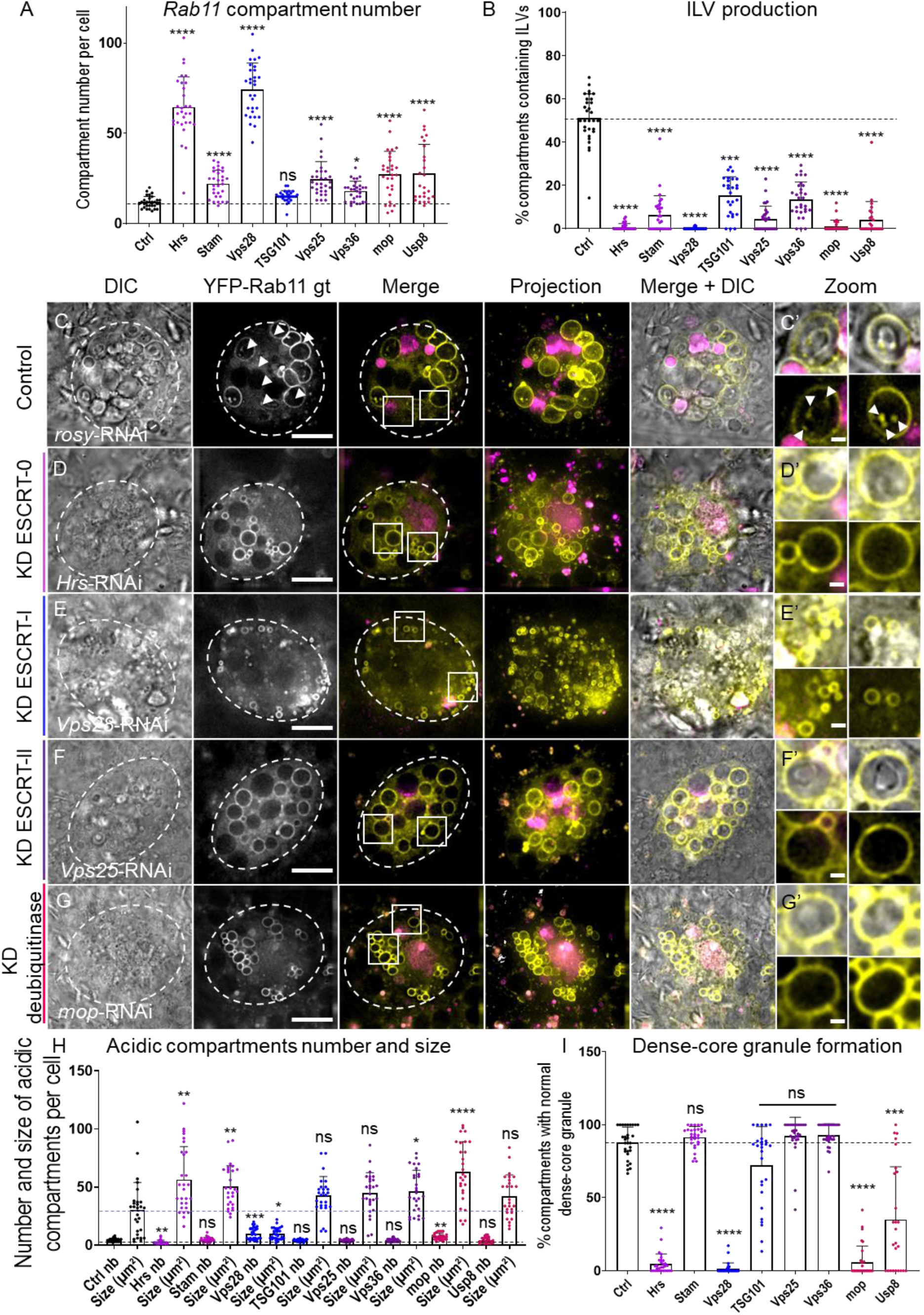
ESCRT 0-II knockdown inhibits exosome biogenesis in non-acidic SC compartments, but does not change their identity. (A) Bar chart showing the number of non-acidic YFP-Rab11-positive compartments with diameter > 0.4 µm per SC in control vs ESCRT-0-II knockdowns. n = 30. (B) Bar chart showing percentage of YFP-Rab11 compartments containing YFP-Rab11-positive ILV puncta. n = 30 SCs. Panels C-G show basal wide-field fluorescence views of living SCs expressing YFP-Rab11 from its endogenous genomic location (yellow) together with *rosy-* or *ESCRT*-RNAis. SC outline approximated by dashed white circles. Acidic compartments are marked by LysoTracker® Red (magenta). Boxed non-acidic compartments are magnified in C’-G’. (C) Control SC expressing *rosy*-RNAi construct. Rab11-positive ILV puncta are present inside 50% of Rab11-compartments (arrowheads; C’). (D) SC expressing *Hrs*-RNAi. Rab11-positive ILVs are absent and size of largest LEL is increased. (E) SC expressing *Vps28*-RNAi. YFP-Rab11-positive ILVs are absent. (F) SC expressing *Vps25*-RNAi. YFP-Rab11-positive ILVs are reduced. (G) SC expressing *mop*-RNAi. YFP-Rab11-positive ILVs are reduced. (H) Bar chart showing number of acidic compartments per SC and maximum area of the largest in µm^2^. n = 30. (I) Bar chart showing proportion of YFP-Rab11 compartments with DCGs in each SC. n = 30. All data are from 6-day-old males shifted to 29°C at eclosion to induce transgene expression of transgenes. Genotypes are: *w; P[w+, tub-GAL80ts]/+; dsx-GAL4/ w; TI{TI}Rab11EYFP* with RNAi #1 lines for each gene. Scale bars in C-G, 10 µm and in C’-G’, 1 µm. Data were analysed by Kruskal-Wallis test. *p < 0.05, **p < 0.01, ***p < 0.001, ****p < 0.0001 relative to control. See also Fig S3 for images of gene knockdowns not depicted here, and Fig S4 for additional RNAi lines tested.

SCs depleted of transcripts encoding the core and accessory ESCRT-0 proteins, Hrs, Stam, Mop and Usp8 had smaller Rab11-compartments than in control cells and a significantly reduced proportion of these contained internal YFP-Rab11 ILVs (Fig 4A,B,D,G, Fig S3B,F and Fig S4A,B). Consistent with our observations with the Btl-GFP marker, DCGs were absent or reduced in size and mis-localised, except with *Stam* knockdown (Fig 4D,G,I and Fig S3B,F). With the exception of *Usp8*-RNAi, all knockdown cells also had enlarged acidic compartments (Fig 4H), as observed in mammalian cells (Razi and Futter, 2006).

Knockdown of the ESCRT-I, *Vps28* (Fig 4E), led to accumulation of many small Rab-11-compartments, which rarely contained YFP-Rab11 puncta or DCGs and severely reduced LEL size, while decreasing *TSG101* levels (Fig S3C) produced a less severe reduction in compartments containing ILVs and no effect on DCGs (Fig 4A,B,H,I, Fig S3C, and Fig S4A, B). When the ESCRT-II genes, *Vps25* and *Vps36*, were knocked down, the resulting SCs had normal acidic compartments and DCGs, but the proportion of compartments with Rab11-positive ILVs was reduced (Figs 4A,B,F,H,I, Fig S3D and Fig S4A,B), again mirroring the effects of these manipulations on Btl-GFP-expressing SCs. In summary, knockdown of ESCRT-0, -I and -II genes does not appear to affect the identity of large non-acidic compartments in SCs, but it does reduce their production of Rab11-labelled ILVs and for several ESCRT-0 and ESCRT-I genes, it also affects DCG formation.

We also investigated how reduction in core and accessory ESCRT-III genes affected SCs. As we had found for Btl-GFP-expressing cells, *shrub* (Fig 5D) and *Vps4* (Fig 5E) knockdowns induced many small Rab11-positive compartments that did not contain puncta or detectable dense cores (Fig 5A, B, I and Fig S4C, D). Late endosomal and lysosomal compartments were also reduced in size (Fig 5H).

**Figure 5.**
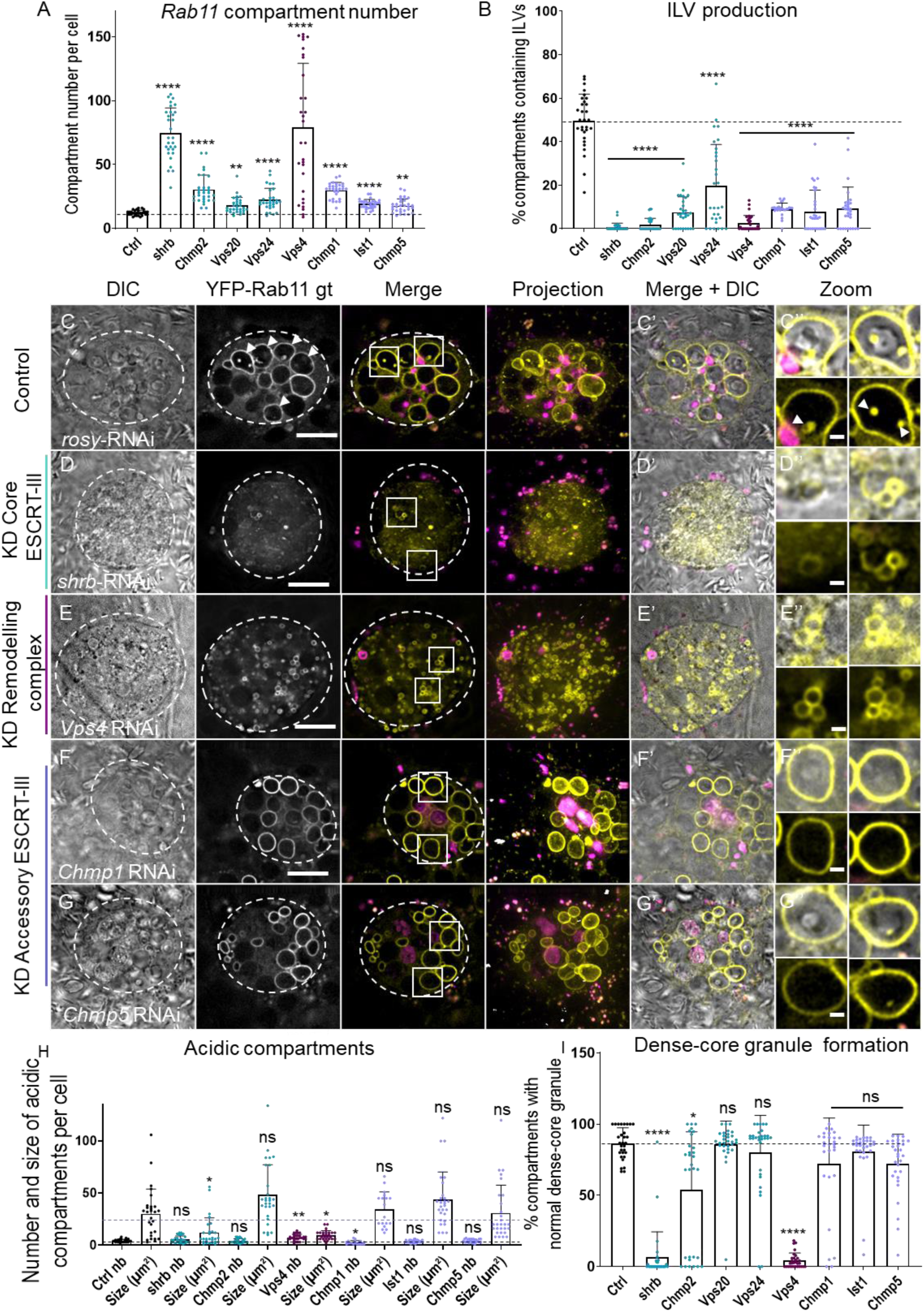
ESCRT III knockdown inhibits exosome biogenesis in non-acidic SC compartments, but does not change their identity. (A) Bar chart showing the number of non-acidic YFP-Rab11-positive compartments with diameter > 0.4 µm per SC in control vs *ESCRT-III* knockdowns. n = 30. (B) Bar chart showing percentage of YFP-Rab11 compartments containing YFP-Rab11-positive ILV puncta. n = 30. Panels C-G show basal wide-field fluorescence views of living SCs expressing YFP-Rab11 from its endogenous genomic location (yellow). SC outline approximated by dashed white circles. Acidic compartments are marked by LysoTracker® Red (magenta). Boxed non-acidic compartments are magnified in C’-G’ Zoom. (C) Control SC expressing *rosy-*RNAi. Rab11-positive ILV puncta are present inside 50% of Rab11-compartments (arrowheads; Zoom). (D) SC expressing *shrub-* RNAi. YFP-Rab11-compartment number is increased, but YFP-Rab11-positive ILVs are strongly reduced. LELs are small or absent. (E) SC expressing *Vps4*-RNAi. YFP-Rab11-positive compartment number is greatly increased. YFP-Rab11-positive ILVs are absent. LELs are numerous and small. (F) SC expressing *Chmp1*-RNAi. YFP-Rab11-positive ILVs are reduced. (G) SC expressing *Chmp5*-RNAi. YFP-Rab11-positive ILVs are reduced. (H) Bar chart showing number of acidic compartments per SC and maximum area of the largest in µm^2^. n = 30. (I) Bar chart showing proportion of YFP-Rab11 compartments with DCGs in each SC. n = 30. All data are from 6-day-old males shifted to 29°C at eclosion to induce transgene expression. Genotypes are: *w; P[w*^*+*^, *tub-GAL80ts]/+; dsx-GAL4/ w; TI{TI}Rab11EYFP* with RNAi #1 lines for each gene. Scale bars in C-G, 10 µm and in C’-G’, 1 µm. Data were analysed by Kruskal-Wallis test. *p < 0.05, **p < 0.01, ***p < 0.001 and ****p < 0.0001 relative to control. See also Fig S4 for images of gene knockdowns not depicted here, and Fig S5 for additional RNAi lines tested.

For knockdown of ESCRT-III core genes *Chmp2, Vps24* and *Vps20*, Rab11-compartments were relatively normal in size, contained DCGs (though they were mildly affected in *Chmp2* knockdown), but only occasionally included YFP-Rab11 puncta (Fig 5A,B,I, Fig S3F-H and Fig S4C,D). These phenotypes were generally milder than those seen with knockdown in Btl-GFP-expressing SCs, although the phenotype of *Chmp2* knockdown cells varied considerably. Acidic compartments were also not significantly affected. Nevertheless, all these genetic manipulations strongly reduced the proportion of non-acidic compartments containing YFP-Rab11 puncta without affecting their identity, essentially mirroring the phenotypes seen with the Btl-GFP marker.

Finally, the three accessory ESCRT-III genes, *Chmp1, Chmp5* and *Ist1* were tested. Knockdown had variable effects on non-acidic compartment size and number. The number of compartments increased in all knockdown SCs, albeit only mildly for Chmp5 and Ist1 knockdown (Fig 5A,F,G Fig S3I and Fig S4C,D). Acidic compartments and DCGs were unaffected (Fig 5H,I), in contrast to some of these manipulations in a Btl-GFP-expressing background (Fig 3). Importantly, knockdown of the accessory ESCRTs consistently reduced the proportion of Rab11-compartments containing Rab11-labelled puncta, indicating that these genes are required for Rab11-exosome biogenesis.

### Accessory ESCRT-III Genes Do Not Affect De-Ubiquitinylation of ILV Cargos in SCs

ESCRTs play a critical role in the degradation of ubiquitinylated proteins in the late endosome by clustering these molecules at the limiting membrane and then sequestering them into ILVs in a process involving deubiquitinylation (Mizuno et al., 2006). We hypothesised that the morphological defects observed in SCs expressing RNAis targeting the core ESCRTs might be partly explained by blockade of this process, which would lead to accumulation of ubiquitinylated ILV cargos. Indeed, in fixed and immunostained AGs from all core ESCRT-0, -I and -III knockdown SCs expressing Btl-GFP, ubiquitin accumulated at readily detectable levels throughout the cell, unlike wild type cells (Fig 6A-H, K and Fig S5A-E). Consistent with the milder defects observed following knockdown of ESCRT-II genes, ubiquitin accumulated to a more limited and non-significant extent in these genetic backgrounds, although for *Vps25* knockdown, almost every gland showed a phenotype that was more severe than all control glands. These results contrasted sharply with knockdowns of accessory ESCRT-III genes, where accumulation of ubiquitin was comparable to wild type cells (Fig 6 I-J, K and Fig S5F). These findings suggest that ESCRT-dependent deubiquitinylation events on late endosomes are not affected by accessory ESCRT-III proteins. They also indicate that the generation of Rab11-exosomes may not involve a ubiquitinylation/deubiquitinylation cycle for cargos, even though it requires ESCRT-0 proteins that are thought to recruit these cargos (Prag et al., 2007).

**Figure 6.**
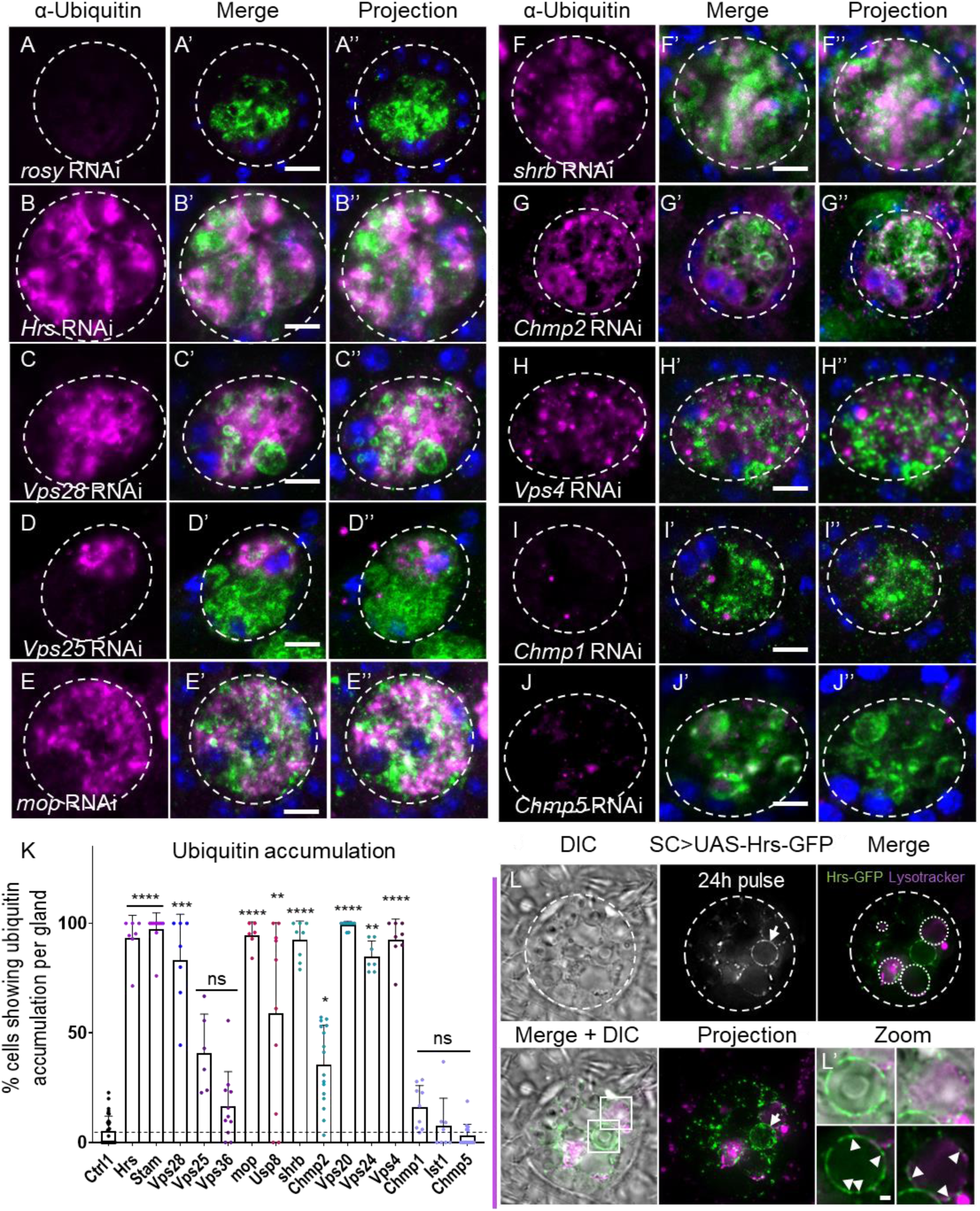
Accessory ESCRT-III proteins are not required for processing of ubiquitinylated ILV cargos. Confocal basal images of fixed SCs isolated from males expressing Btl-GFP and selected ESCRT-RNAis from eclosion onwards. SC outline approximated by dashed white circles. Ubiquitin (magenta), GFP (green) and DAPI (blue; marking binucleate cells) staining is shown. Nuclear staining with anti-ubiquitin is sometimes observed, even in controls, but appears non-specific. (A) Control SC expressing *rosy*-RNAi. Virtually no cytoplasmic accumulation of ubiquitin is observed. (B) SC expressing *Hrs*-RNAi reveals strong accumulation of ubiquitin in cytosol and some Btl-GFP-positive compartments (colocalization in white). (C) SC expressing *Vps28*-RNAi accumulates high levels of ubiquitin. (D) SC expressing *Vps25*-RNAi shows a milder accumulation of ubiquitin. (E) SC expressing *mop*-RNAi accumulates ubiquitin, including in Btl-GFP-positive compartments. (F) SC expressing *shrub*-RNAi accumulates ubiquitin in the cytosol and Btl-GFP-positive compartments. (G) SC expressing *Chmp2*-RNAi accumulates ubiquitin in the cytosol and Btl-GFP-positive compartments. (H) SC expressing *Vps4*-RNAi accumulates ubiquitin in the cytoplasm. (I) SC expressing *Chmp1*-RNAi does not accumulate ubiquitin in the cytoplasm. (J) SC expressing *Chmp5*-RNAi does not accumulate ubiquitin in the cytoplasm. (K) Bar chart showing proportion of SCs per gland that accumulate ubiquitin in the cytoplasm in control and ESCRT knockdowns. n < 6AGs. (L) Basal view through a living SC from a 6-day-old male, following a 24 h pulse of *UAS-Hrs-GFP* transgene expression at 29°C. In addition to punctate localisation at the surface of LELs (dotted white circles in top right image and arrowheads in bottom right-hand images in Zoom), which are marked by the vital dye LysoTracker® Red (magenta), Hrs-GFP (green) is found in foci (arrowheads in left-hand images in Zoom) at the membrane of some non-acidic compartments (arrow). Images were acquired by wide-field microscopy. Genotype is *w; P[w*^*+*^, *tub-GAL80ts]/+; dsx-GAL4/P[w*^*+*^, *UAS-Hrs-GFP]*. All data, except L, are from 6-day-old male flies shifted to 29°C at eclosion to induce transgene expression. Genotypes are: *w; P[w*^*+*^, *tub-GAL80ts]/+; dsx-GAL4/P[w*^*+*^, *UAS-btl-GFP]* with *rosy-* or RNAi #1 lines. Scale bars in A-L, 10 µm. Data were analysed by Kruskal-Wallis test. *p < 0.05 **p < 0.01 ***p < 0.001 ****p < 0.0001 relative to control. See Fig S5 for images of gene knockdowns not depicted here.

We previously showed that the core ESCRT-III protein Shrub is present in subdomains on the limiting membrane of Rab11-compartments (Fan et al., 2019). To test whether the ESCRT-0 complex might also concentrate around these compartments, as well as LELs, GFP-tagged Hrs was expressed over a 24 h period in SCs. It localised to regions on the limiting membrane of at least one DCG-containing compartment in most SCs (10/14 cells), in addition to marking foci on the LEL limiting membrane, consistent with the ESCRT-0 complex playing a direct role in Rab11-exosome biogenesis (Fig 6L).

We have previously used a GFP-tagged form of the putative human exosome marker CD63 (Escola et al., 1998; Pols and Klumperman, 2009) to study Rab11-exosome biogenesis and endosomal trafficking in SCs (Corrigan et al., 2014; Fan et al., 2019). CD63-GFP appears to enhance endosomal trafficking; it marks Rab11-exosomes when overexpressed, and also traffics to enlarged LELs in these cells, where internalised GFP is quenched by the acidic environment (Corrigan et al., 2014). To confirm our findings with Btl-GFP, we tested whether ubiquitin failed to accumulate in CD63-GFP-expressing SCs following knockdown of accessory ESCRT-III genes. Screening a subset of *ESCRT* knockdowns revealed similar phenotypes to those seen with Btl-GFP: a general decrease in ILV formation and exosome secretion for all knockdowns (Fig 7). Co-staining for GFP and ubiquitin highlighted ubiquitin accumulation following core *ESCRT* knockdowns (Fig 7B’’-E’’), but not when accessory ESCRT-III expression was reduced (Fig 7F’’, I). We conclude that although accessory ESCRT-IIIs are required for Rab11-exosome biogenesis, they do not appear to be essential for the processing of ubiquitinylated cargos during ILV formation in late endosomes.

**Figure 7.**
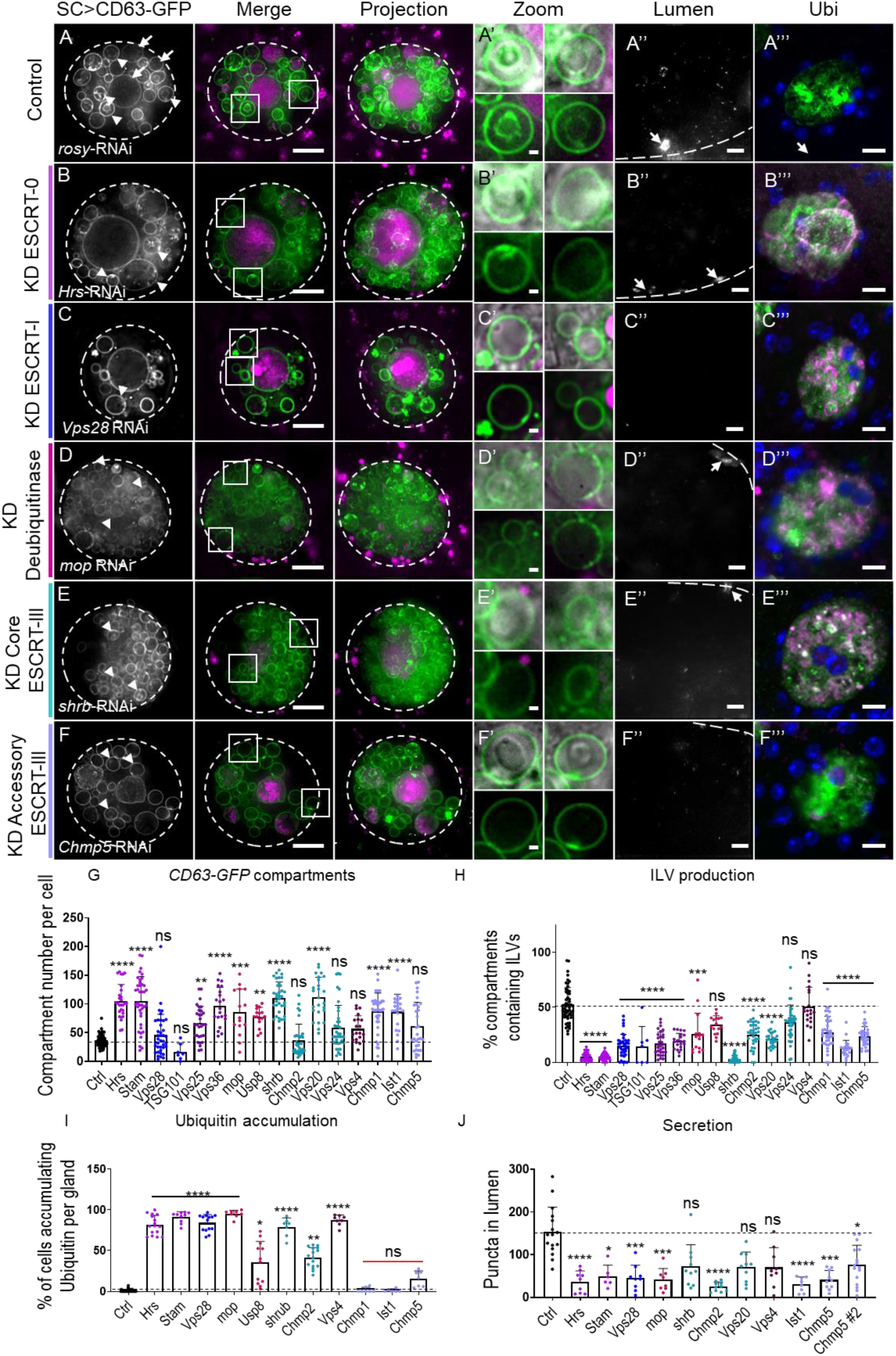
Knock down of accessory ESCRT-III components suppresses Rab11-exosome formation but not processing of ubiquitinylated ILV cargos. Panels A-F show basal wide-field fluorescence views of living SCs expressing *UAS-CD63-GFP* (green). SC outline approximated by dashed white circles. Acidic compartments are marked by LysoTracker® Red (magenta). CD63-GFP labels most large endosomal compartments in SCs and ILVs within them, both acidic (arrows; shown in control) and non-acidic (arrowheads). Boxed non-acidic compartments are magnified in A’-F’. Confocal transverse images of AG lumens are shown in A’’-F’’; SCs are highlighted with arrows and luminal exosomes appear as puncta. A’’’-F’’’ show confocal basal images of fixed SCs with staining for GFP (green), ubiquitin (magenta) and DAPI (blue; cells in AG are binucleate). (A) Control SC expressing *rosy*-RNAi. CD63-GFP-positive ILVs are found around DCGs and in clusters that link to the limiting membrane of non-acidic compartments (A’). Exosomes are also visible as puncta in the AGlumen (A’’). Ubiquitin does not accumulate in the SC cytoplasm (A’’’). (B) SC expressing *Hrs*-RNAi. Largest lysosome (magenta) is enlarged (Merge) and CD63-GFP-positive ILVs in non-acidic compartments (B’) and exosome secretion are reduced (B’’). Ubiquitin accumulates in cytoplasm (B’’’). (C) SC expressing *Vps28*-RNAi. CD63-GFP-positive ILVs in non-acidic compartments (C’) and exosome secretion (C’’) are reduced. Ubiquitin accumulates in cytoplasm (C’’’). (D) SC expressing *mop*-RNAi. CD63-GFP-positive ILVs in non-acidic compartments (D’) and exosome secretion (D’’) are reduced. Ubiquitin accumulates in the cytoplasm (D’’’). (E) SC expressing *shrub*-RNAi. CD63-GFP-positive ILVs in non-acidic compartments (E’) and exosome secretion (E’’) are reduced. Ubiquitin accumulates in the cytoplasm (E’’’). (F) SC expressing *Chmp5*-RNAi. CD63-GFP-positive ILVs are strongly reduced in non-acidic compartments (F’). Secretion is reduced (F’’). Ubiquitin does not accumulate in the cytoplasm (F’’’). (G) Bar chart showing the number of non-acidic CD63-GFP-positive compartments with diameter > 0.4 µm per SC in control vs *ESCRT* knockdowns. n = 30. (H) Bar chart showing percentage of CD63-GFP-positive compartments containing GFP-positive ILVs. n = 30. (I) Bar chart showing proportion of cells per gland that accumulate ubiquitin in the cytoplasm of SCs. n ≥ 6. (J) Bar chart of number of CD63-GFP-positive puncta in AG lumen. n ≥ 6. All data are from 6-day-old male flies shifted to 29°C at eclosion to induce transgene expression. Genotypes are: *w; P[w*^*+*^, *tub-GAL80ts]/+; dsx-GAL4/P[w*^*+*^, *UAS-CD63-GFP]* with *UAS-rosy-RNAi* or *UAS-ESCRT*-*RNAi*-#1 constructs for *Hrs, mop, Usp8, Vps28, Vps36, shrb, Vps20, Chmp2, Vps4, Chmp1, Ist1, Chmp5* and -#3 for *Stam, TSG101, Vps25, Vps24*. Scale bars, 10 µm, except A’-F’, 1 µm. Data were analysed by Kruskal-Wallis test. *p < 0.05, **p < 0.01, ***p < 0.001 and ****p < 0.0001 relative to control.

## Discussion

Rab11-positive recycling endosomes have recently been identified as a source of exosomes, in addition to the well-established late endosomal pathway. Exosomes from the former compartments carry different cargos and in cancer cells, appear to have novel functions that may be involved in adaptation. In *Drosophila*, some of the core ESCRTs have previously been reported to control Rab11-exosome biogenesis (Fan et al., 2019), mirroring their roles in forming late endosomal exosomes (Schuh and Audhya 2014). Here we present data supporting a more selective role for accessory ESCRT-III proteins in Rab11-exosome production and propose that these proteins promote ILV formation via an ESCRT-dependent mechanism distinct from late endosomes.

### Core ESCRTs control Rab11- and late endosomal exosome biogenesis

All four ESCRT complexes have previously been implicated in the formation of multivesicular endosomes in yeast; *ESCRT* mutants produce endosomes without ILVs and so-called ‘class E’ endosomal compartments with abnormal tubulation (Raymond et al., 1992; Rieder et al., 1996; Babst et al., 2002a). In higher organisms, there is evidence for ESCRT-independent exosome production (Trajkovic et al., 2008); it has also been reported that cells depleted of components from any of the four core ESCRT complexes can generate CD63-positive MVEs in metazoa via a ceramide-dependent mechanism (Stuffers et al., 2009). However, exosome secretion from multivesicular late endosomes is heavily dependent on ESCRTs in multiple cell types (Möbius et al., 2002; White et al., 2006; Colombo et al., 2014), permitting secretion of ILV-associated receptors, ligands and cytosolic cargos with a wide range of signalling roles (Matusek et al., 2014). In this study, we have shown that ESCRT proteins from all ESCRT complexes are needed for normal Rab11-exosome biogenesis in fly SCs, and that most of them also control the proper size and morphology of large SC secretory compartments, but not their Rab11 identity. Previous reports have demonstrated that ESCRT-0 *Hrs* mutants induce the expansion of acidic endosomes (Razi and Futter, 2006). *Hrs, Stam* and *mop* knockdown all increase the size of the largest acidic compartment in SCs (Fig 4H), suggesting there are other parallels in ESCRT function between SCs and mammalian cells.

Core *ESCRT-II* knockdowns produced the most variable phenotypes. The proportion of ILV-containing non-acidic compartments was partially reduced in SCs expressing all three markers (Figs 2, 4, 7, S1-S4), but in Btl-GFP-expressing cells, the changes were modest and frequently accompanied by the formation of large ILVs in some compartments. The effects on ubiquitin accumulation were also milder than with other core *ESCRT* knockdowns (Fig 6). It is unclear whether this can simply be explained by a weaker knockdown of these genes in SCs or is the result of a mechanistic difference. Interestingly in yeast, Doa4 (the deubiquitinase for ILV cargo in S. cerevisiae) interacts specifically with ESCRT-III proteins Vps20, Vps24 and Snf7 (yeast Shrb), but does not bind to the ESCRT-II Vps25 (Wolters and Amerik, 2015; Johnson et al., 2017). In flies, Usp8, which produces a strong ubiquitin accumulation phenotype following knockdown (Fig S5D), appears to be the functional orthologue of Doa4, and might also partially bypass ESCRT-II components in SCs in the cycling of modified cargos during ILV formation.

### Dense-core granule formation and Rab11-exosome biogenesis appear to be functionally associated

Many of the more severe *ESCRT* knockdown phenotypes were not only characterised by a reduction in ILV-forming compartments, but also by effects on DCGs in non-acidic compartments. This was most clearly observed in SCs expressing Btl-GFP (Fig 2H), where knockdowns of transcripts encoding ESCRT-0 components (other than Stam), the ESCRT-I Vps28, and several core ESCRT-III proteins, either appeared to block DCG formation entirely (eg. Fig 2E,G) or in some compartments, lead to formation of one or more small mis-localised DCGs (eg. Fig 2D,F,G, Fig 3B,E and Fig S1B,G). Although absence of DCGs might be partly associated with the reduced size of non-acidic compartments in these genetic backgrounds, leading to small cores that cannot be observed using DIC microscopy (eg. Fig 2D, E Fig 3B-E), these phenotypes were observed even in larger compartments. In knockdown cells expressing the *YFP-Rab11* gene trap, defects were milder, with DCGs being small and/or fragmented (Fig 4D,E,G, Fig 5D,E, and Fig S3E) in knockdowns with the most severe phenotypes.

Currently we do not understand how ESCRT function and DCG biogenesis are linked, although it seems unlikely DCG formation is simply dependent on ILVs, because *Stam1* and *TSG101* knockdowns strongly suppresses ILV formation, but leave DCG number unaffected. Interestingly, we recently reported a related phenotype in SCs with reduced expression of the glycolytic enzyme, glyceraldehyde dehydrogenase 2 (GAPDH2; (Dar et al., 2020)). Multiple small dense cores, surrounded by ILVs, form in close proximity to the limiting membrane of compartments. Since extravesicular GAPDH2 appears to promote ILV clustering, one possible explanation for this phenotype is that clusters of ILVs in Rab11-compartments, which normally surround each core and link it to the limiting membrane, are important for assembly of the large dense cores in SCs. Perhaps only some ESCRTs contribute to the events leading to this clustering process. Whether the same mechanisms are used in other cells with much smaller dense-core vesicles remains unclear.

### Accessory ESCRT-III knockdown inhibits Rab11-exosome biogenesis in secondary cells, but ubiquitinylated endosomal cargos do not accumulate

We have found that accessory ESCRT-III proteins are required for Rab11-exosome biogenesis (Fig5, Fig S3). These molecules have previously been postulated to be modulators of the vesicle scission activities of core ESCRT-III components (Azmi et al., 2008; Rue et al., 2007). The Chmp1-Ist1 subcomplex is thought to facilitate the recruitment of Vps4 to ESCRT-III (Rue et al., 2007; Agromayor et al., 2009) and the Chmp5-Vta1 complex to enhance Vps4 activity (Shiflett et al., 2004; Azmi et al., 2006, 2008; Xiao et al., 2008). In the majority of studies, including in *Drosophila*, a single accessory ESCRT-III knockdown or mutation appears to have either a subtle or no phenotype (Vaccari et al., 2009), and loss of an additional accessory ESCRT-III protein is required to generate a strong phenotype (Bäumers et al., 2019).

In mammals, loss of *Chmp5* function is associated with early embryonic lethality (Shim et al., 2006). In *chmp5* ^*-/-*^ primary embryonic cell cultures, enlarged late endosomes packed with vesicles are observed, suggesting that ILVs can still be made under these conditions. *Lip5/Vta1* and *Chmp5* knockdowns in HeLa cells lead to EGFR accumulation (Ward et al., 2005), suggesting that aspects of endosomal trafficking are defective, but effects on ubiquitinylation and ILV formation were not investigated.

In one example that parallels our findings, the accessory ESCRT-III Ist1 is required to maintain cargo clustering and ILV formation in the *C. elegans* oocyte-to-embryo transition (Frankel et al., 2017). At this stage, MVE biogenesis appears rapid, as it is in SCs. It will be interesting to investigate whether these MVEs are late endosomal or recycling endosomal in origin.

Although accessory *ESCRT-III* knockdowns block Rab11-exosome biogenesis, we observed two major differences compared to knockdowns of most core ESCRTs in SCs. First, the size of Rab11 compartments and their morphology was less affected by reduction of accessory ESCRT-III proteins. Second, there was no obvious accumulation of ubiquitin. Ubiquitin accumulation following core *ESCRT* knockdown is thought to reflect the involvement of these ESCRTs in clustering ubiquitinylated cargos on the late endosomal limiting membrane and then deubiquitinylating them as ILVs form. Our accessory *ESCRT-III* knockdown data suggest either that this ubiquitinylation/deubiquitinylation cycle can take place even in the absence of ILV formation, or perhaps more likely, that the cycle is not involved in Rab11-exosome biogenesis and late endosomal exosomes continue to be generated in these knockdown cells.

In this regard, it is interesting to note that others have reported ESCRT-dependent ILV biogenesis events that do not require ubiquitinylation. Most notably, stress-internalised EGFR is trafficked to an alternative form of MVEs in human cells that is ESCRT-dependent, but does not require ubiquitin for EGFR trafficking into ILVs (Tomas and Futter, 2015).

The implication of core ESCRTs, like ESCRT-0 Hrs, which is thought to interact with ILV cargos via its ubiquitin-interacting motif in ubiquitin-independent ILV formation is intriguing. For example, ubiquitinylated Transferrin Receptors (TfRs) interact with Hrs and are sorted to the degradative pathway, whereas non-ubiquitinylated endocytosed receptors fail to colocalize with Hrs and rapidly recycle to the cell surface (Raiborg et al., 2001; Matusek et al., 2019). Since Rab11-exosomes are generated in recycling compartments, could their cargos be recruited via a different mechanism, for example, via a ubiquitin-like protein (Hochstrasser, 2009). In this regard, human Chmp4B (equivalent to *Drosophila* Shrub), Chmp1A and Chmp5 have been shown to directly interact with the SUMO-conjugating enzyme Ubc9 (Tsang et al., 2006). Therefore, sumoylation is one process that requires further investigation, particularly because it has been linked to recruitment of some cargos into exosomes (Villarroya-Beltri et al., 2013).

Since pro-tumorigenic Rab11a-exosomes are secreted under nutrient stress conditions in cancer cell lines (Fan et al., 2019), our data could be relevant to the study of tumour adaptation mechanisms. Some accessory ESCRT-IIIs have already been associated with human cancer: Ist1 overexpression (OLC1 in human also known as KIAA0174) has been observed in lung, breast and colorectal cancer as well as oesophageal squamous cell carcinoma (Yuan et al., 2008; Ou-Yang et al., 2014; Liu et al., 2014; Li et al., 2014), and Chmp1A levels are reported to be increased in renal cell and pancreatic carcinomas (Li et al., 2008; Mochida et al., 2012, p. 1). It may be possible to selectively inhibit Rab11a-exosome biogenesis by interfering with accessory ESCRT-III function and therefore block adaptive responses without preventing the ubiquitin-mediated degradation of growth factor receptors in cancer cells.

Overall, we conclude that accessory ESCRT-III proteins selectively regulate Rab11-exosome formation in SCs via an ESCRT-dependent mechanism, which does not appear to be dependent on a ubiquitinylation/deubiquitinylation cycle for cargo loading. Further investigation of this novel loading mechanism should begin to unravel how different exosome subtypes are made, and how this affects exosome function under different cellular conditions.

## Materials and methods

### Drosophila Stocks and Genetics

The following fly stocks, acquired from the Bloomington *Drosophila* Stock Centre (RNAi lines from the TRiP collection; (Ni et al., 2009) and the Vienna Drosophila Research Center (RNAi lines from the shRNA, GD and KK libraries; (Dietzl et al., 2007), unless otherwise stated, were used: *UAS-rosy-RNAi-*#1 (TRiP.HMS02827, BL 44106), *UAS-Hrs-RNAi-*#1 (TRiP.JF02860, BL 28026) (Sheng et al., 2016), *-*#2 (TRiP.HMS00841, BL 33900) (Gomez-Lamarca et al., 2015; Loncle et al., 2015), UAS-*Stam-RNAi-*#1 (GD 11948, VDRC 22497) (Bras et al., 2012; Gomez-Lamarca et al., 2015), *-*#2 (TRiP.HMS01429, BL 35016) (Sheng et al., 2016), -#3 (shRNA 330248, VDRC 330248), *UAS-Mop*-RNAi*-*#1 (TRiP.HMS00706, BL 32916) (Loncle et al., 2015), -#2 (TRiP.HM05008, BL 28522), *UAS-Usp8*-RNAi*-*#1 (TRiP.HMS01941, BL 39022), -#2 (TRiP.HMS01898, BL 38982) (Loncle et al., 2015), *UAS-TSG101*-RNAi*-*#1 (TRiP.GLV21075, BL 35710) (Loncle et al., 2015), -#2 (GD 14295, VDRC 23944) (Bras et al., 2012; Mamińska et al., 2016), *UAS-Vps28*-RNAi*-*#1 (GD 7696, VDRC 31894) (Bras et al., 2012; Mamińska et al., 2016; Neyen et al., 2016), -#2 (KK 101474, VDRC 105124) (Mamińska et al., 2016), *UAS-Vps36*-RNAi*-*#1 (TRiP.HMS01739, BL 38286) (Loncle et al., 2015), -#2 (KK 102099, VDRC 107417), UAS-Vps25 -RNAi*-*#1 (GD 8432, VDRC 38821) (Bras et al., 2012), -#2 (KK 102944, VDRC 108105) (Neyen et al., 2016), -#3 (TRiP.JF02055, BL 26286) (Loncle et al., 2015), *UAS-shrb-RNAi-*#1 (KK 108557, VDRC 106823) (Bras et al., 2012), -#2 (TRiP.HMS01767, BL 38305), *UAS-Vps24-RNAi-*#1 (GD 14676, VDRC 29275), -#2 (TRiP.HMS01733, BL 38281) (Loncle et al., 2015), -#3 (KK 107601, VDRC 100295), *UAS-Vps20-RNAi-*#1 (GD 11211, VDRC 26387) (Bras et al., 2012), -#2 (discontinued, VDRC 47653), UAS-*chmp2-RNAi-*#1 (GD 8363, VDRC 24869) (Bras et al., 2012), -#2 (TRiP.HMS01911, BL 38995) (Loncle et al., 2015), *UAS-Chmp1-RNAi-*#1 (GD 11219, VDRC 21788) (Bras et al., 2012), -#2 (TRiP.HM05117, BL 28906) (Bras et al., 2012), *UAS-Ist1-RNAi-* #1 (GD 6866, VDRC 31174), -#2 (KK 108546, VDRC 100771), *UAS-Chmp5-RNAi-*#1 (KK 109120, VDRC 101422) (Loncle et al., 2015), -#2 (GD 10565, VDRC 25990) (Bras et al., 2012), *UAS-Vps4-RNAi-*#1 (KK 101722, VDRC 105977) (Bras et al., 2012), -#2 (GD 12054, VDRC 35125) (Bras et al., 2012). Most of these lines have been validated by independent screens (Bras et al., 2012; Gomez-Lamarca et al., 2015; Loncle et al., 2015; Mamińska et al., 2016; Neyen et al., 2016; Sheng et al., 2016). Appropriate genotypes of both sexes were used in this study.

Flies were grown at 25°C in vials containing standard cornmeal agar medium (per litre, 12.5 g agar, 75 g cornmeal, 93 g glucose, 31.5 g inactivated yeast, 8.6 g potassium sodium tartrate, 0.7 g calcium, and 2.5 g Nipagen dissolved in 12 ml ethanol). They were transferred onto fresh food every 3-4 days. No additional dried yeast was added to the vials. The specific driver *dsx-GAL4* and the temperature-sensitive, ubiquitously expressed, repressor *tubulin-GAL80*^*ts*^ were used to induce SC-specific expression of *UAS-CD63-GFP* or *UAS-Btl-GFP* in a temperature-controlled fashion. Expression of the *YFP-Rab11* gene trap was controlled by *Rab11* regulatory sequences. Females carrying *dsx-GAL4, tubulin-GAL80*^*ts*^ and one of the three compartment markers were crossed with males carrying the different UAS-RNAi transgenes. Virgin male offspring, isolated after eclosion, were transferred to 29°C for 6 days to induce post-differentiation SC-specific expression of inducible markers and RNAi.

### Preparation of Accessory Glands for Live Imaging

Accessory glands were prepared following the protocol described by (Fan et al., 2019). In brief, adult male flies were anaesthetised using CO_2_ and dissected in ice-cold 1× PBS (Gibco, Thermo Fisher Scientific). The whole male reproductive system was carefully pulled out of the body cavity, keeping testis, accessory gland, ejaculatory duct and ejaculatory bulb to avoid stress in the tissue. They were then stained with ice-cold 500 nM LysoTracker® Red DND-99 (Invitrogen®) in 1 × PBS for 5 min, washed in ice-cold 1 × PBS for 1 min, then mounted onto High Precision microscope cover glasses (thickness No. 1.5H, Marienfeld-Superior). A custom-built holder was used to hold the specimen during imaging. To avoid dehydration and hypoxia, the glands were maintained in a small drop of 1 × PBS, surrounded by 10S Voltalef® (VWR Chemicals), held by a small cover glass (VWR).

### Microscopy

Live imaging followed the method described by Fan et al. (2019) using the inverted wide-field fluorescence system DeltaVision Core from Olympus AppliedPrecision with a EM-CCD camera, capable of fluorescence and interference contrast (DIC) microscopy. Images and Z-sections for quantification of intracellular compartments were captured with 0.3 µm spacing using the SoftWoRx software.

Fixed tissues were imaged with an upright laser scanning confocal Zeiss LSM880. Zeiss Plan Apochromat oil differential interference contrast objective 63×/1.4 was used with RI 1.514 immersion oil (Zeiss). Z-sections and images for quantification of intracellular compartments were captured with 0.5 µm spacing using the Zen blue suite software according to previously published protocols (Corrigan et al., 2014).

### SC Compartment Analysis

Compartments were analysed and ILV-containing compartments counted following the methods described in Fan et al. (2019), except all compartment with diameter ≥ 0.4 µm were included. Three different SCs were analysed from each of 10 glands. Dense-cores were detected by DIC. If the DCG in a compartment was broken, smaller than control or absent, the compartment was scored as abnormal. For each SC, % of normal DCGs was quantified.

### Analysis of ubiquitin accumulation in SCs

Glands were dissected from virgin six-day-old controls and males expressing *ESCRT*-RNAis in their SCs. They were fixed in 4% paraformaldehyde dissolved in PBS for 15 minutes to preserve tissue integrity and washed in PBST (PBS and 0.3% Triton X-100) for 5 min. Glands were then pre-incubated with 10% goat serum in PBST for one hour and then incubated overnight with primary antibody against GFP (rabbit monoclonal; Abcam, 1:500) and conjugated ubiquitin (Ubi FK2 mouse monoclonal; Millipore, 1:500) in PBST. Glands were rinsed 5 times for 10 minutes in PBST and incubated overnight in PBST with secondary antibody (Alexa555 anti-mouse, 1:400 and Alexa448 anti-rabbit, 1:400; Abcam), then rinsed 5 more times with PBS. Finally, glands were mounted on a glass slide and immersed in a drop of Vectashield with DAPI (Vector Laboratories) and covered with a coverslip for imaging by confocal microscopy.

### *Drosophila* Exosome Secretion Assay

Virgin six-day-old SC>Btl-GFP- or SC>CD63-GFP-expressing males were dissected in ice-cold PBS. Exosome secretion was measured by acquiring a 10 µm z stack within the central third of each gland, with each image spaced by 0.3 µm, using the DeltaVision Elite wide-field microscope. Fluorescence coming from nearby SCs interferes with the acquisition in wide-field microscopy. The automated analysis of exosome secretions by SCs was performed using ImageJ2 (Schindelin et al., 2015), distributed by Fiji. A threshold was set for the raw data from each lumenal area using the Kapur-Sahoo-Wong (Maximum Entropy) method (Kapur et al., 1985) to determine the outlines of fluorescent particles. The ‘Analyse Particles’ function was then used to determine the total number of fluorescent puncta.

### Statistical Analysis

In all analyses, mean values per cell are presented for pooled data and errors bars are SD. For comparison between control and RNAi lines, statistical significance was determined using multi comparisons with one-way ANOVA test. Most data for compartment number counts, and % of compartments producing ILVs or DCGs failed the Shapiro-Wilk normality test (p < 0.01). As a result, the nonparametric Kruskal-Wallis test was used to test significance of differences under all knockdown conditions. Statistical analyses were performed using GraphPad Prism version 8. 3. 1. *p < 0.05, **p < 0.01, ***p < 0.001 and ****p < 0.0001 relative to control.

## Acknowledgements

We are grateful to N. Halidi, R. Parton, D. Pinto and all the staff at the Wellcome Trust-funded MICRON Oxford Advanced Bioimaging Unit, where imaging was undertaken. We thank S. Eaton, S. Goodwin, E. Prince and F. Karch, as well as the Bloomington and Vienna and Stock Centres for *Drosophila* stocks. We acknowledge the support of Cancer Research UK (C19591/A19076, C602/A18974), the BBSRC (BB/K017462/1, BB/N016300/1, BB/R004862/1), and the Wellcome Trust (Strategic Awards #091911, #107457).

## Supplementary figures

**Supplementary Figure S1.**
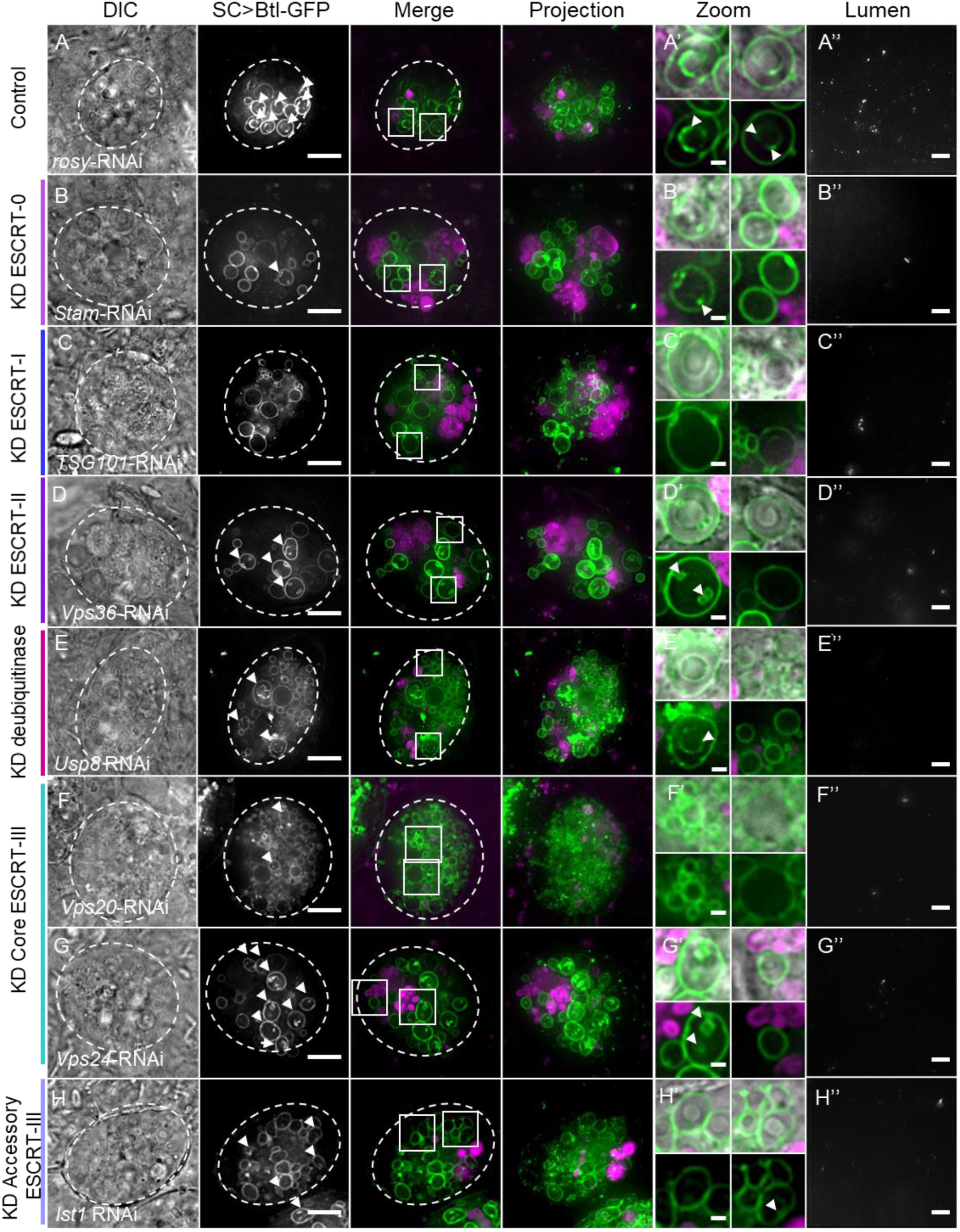
ESCRTs regulate exosome biogenesis in non-acidic compartments of *Drosophila* SCs. Panels A-H show basal wide-field fluorescence views of living SCs from 6-day old males expressing the GFP-tagged form of Breathless (Btl-GFP; green) and a selected RNAi from eclosion onwards. SC outline approximated by dashed white circles. Acidic compartments are marked by LysoTracker® Red (magenta). Boxed non-acidic compartments are magnified in A’-H’. (A) Control SC expressing *rosy*-RNAi construct. Btl-GFP-positive ILVs are visible in many non-acidic compartments (arrowheads; A’) and as puncta in the projection of the AG lumen (A’’). (B) SC expressing *Stam*-RNAi. Acidic compartment size is increased. Btl-GFP-positive ILVs and puncta in the AG lumen are reduced, but DCGs are still present (B’). (C) SC expressing *TSG101*-RNAi has less Btl-GFP-positive ILVs and secreted puncta. (D) SC expressing *Vps36*-RNAi has a lower proportion of non-acidic compartments containing Btl-GFP-positive puncta and ILVs are often enlarged and exosome secretion is reduced. (E) SC expressing *Usp8-*RNAi has less Btl-GFP-positive ILVs and DCGs, and exosome secretion is reduced. (F) SC expressing *Vps20*-RNAi appears to have reduced Btl-GFP-positive ILVs and secreted puncta, though the latter is not significant. (G) SC expressing *Vps24*-RNAi has less Btl-GFP-positive ILVs and secreted puncta. (H) SC expressing *Ist1*-RNAi has normal numbers of Btl-GFP-positive compartments, but less contain ILVs and secreted puncta are reduced. Genotypes are: *w; P[w*^*+*^, *tub-GAL80ts]/+; dsx-GAL4/P[w*^*+*^, *UAS-btl-GFP]* with *UAS-rosy-RNAi* (A), *UAS-Stam-RNAi*-#1 (B), *UAS-TSG101-RNAi-#*1 (C), *UAS-Vps36-RNAi*-#1 (D),*UAS-Usp8-RNAi*-#1 (E), *UAS-Vps20-RNAi*-#1 (F) *UAS-Vps24-RNAi*-#1 (G), and *UAS-Ist1-RNAi*-#1 (H). Scale bars in A-H and AG lumen, 10 µm and in A’-H’, 1 µm.

**Supplementary Figure S2.**
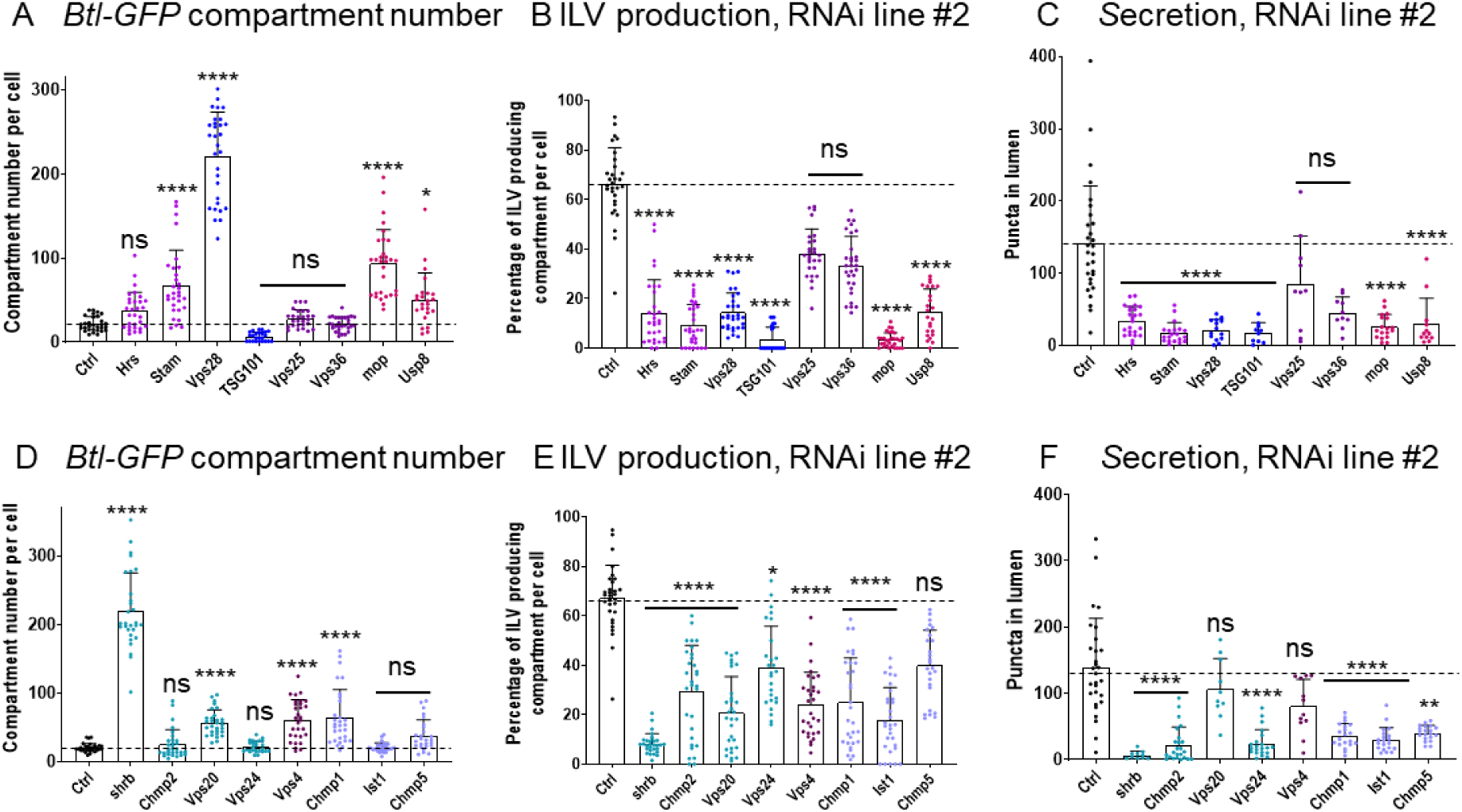
ESCRTs regulate exosome biogenesis in non-acidic compartments of SCs. Quantification of live SCs expressing a second RNAi line (#2) for all ESCRT genes shown in Figs 1, 2 and S1. (A) Bar chart showing the number of non-acidic Btl-GFP-positive compartments with diameter > 0.4 µm per SC in control vs ESCRT-0, -I, -II and deubiquitinases *mop* and *Usp8* knockdowns. n = 30. (B) Bar chart showing percentage of Btl-GFP-positive compartments containing fluorescent ILVs for ESCRT-0, -I, -II and deubiquitinases *mop* and *Usp8* knockdowns. n = 30. (C) Bar chart showing number of Btl-GFP fluorescent puncta in AG lumen following ESCRT-0, -I, -II and deubiquitinases *mop* and *Usp8* knockdown compared to control SCs. n > 10. (D) Bar chart showing the number of non-acidic Btl-GFP-positive compartments with diameter > 0.4 µm per SC in control vs *ESCRT-III* and *Vps4* knockdowns. n = 30. (E) Bar chart showing percentage of Btl-GFP-positive compartments containing fluorescent ILVs for *ESCRT-III* and *Vps4* knockdowns. n = 30. (F) Bar chart showing number of Btl-GFP fluorescent puncta in AG lumen following *ESCRT-III* and *Vps4* knockdown compared to control SCs. n > 10. All data are from 6-day-old males shifted to 29°C at eclosion to induce expression of transgenes. Genotypes are: *w; P[w*^*+*^, *tub-GAL80ts]/+; dsx-GAL4/P[w*^*+*^, *UAS-btl-GFP]* with *UAS-rosy-RNAi*, and RNAi #2 lines for each *ESCRT*. Data were analysed by Kruskal-Wallis test. *p < 0.05, **p < 0.01, ***p < 0.001, and ****p < 0.0001 relative to control.

**Supplementary Figure S3.**
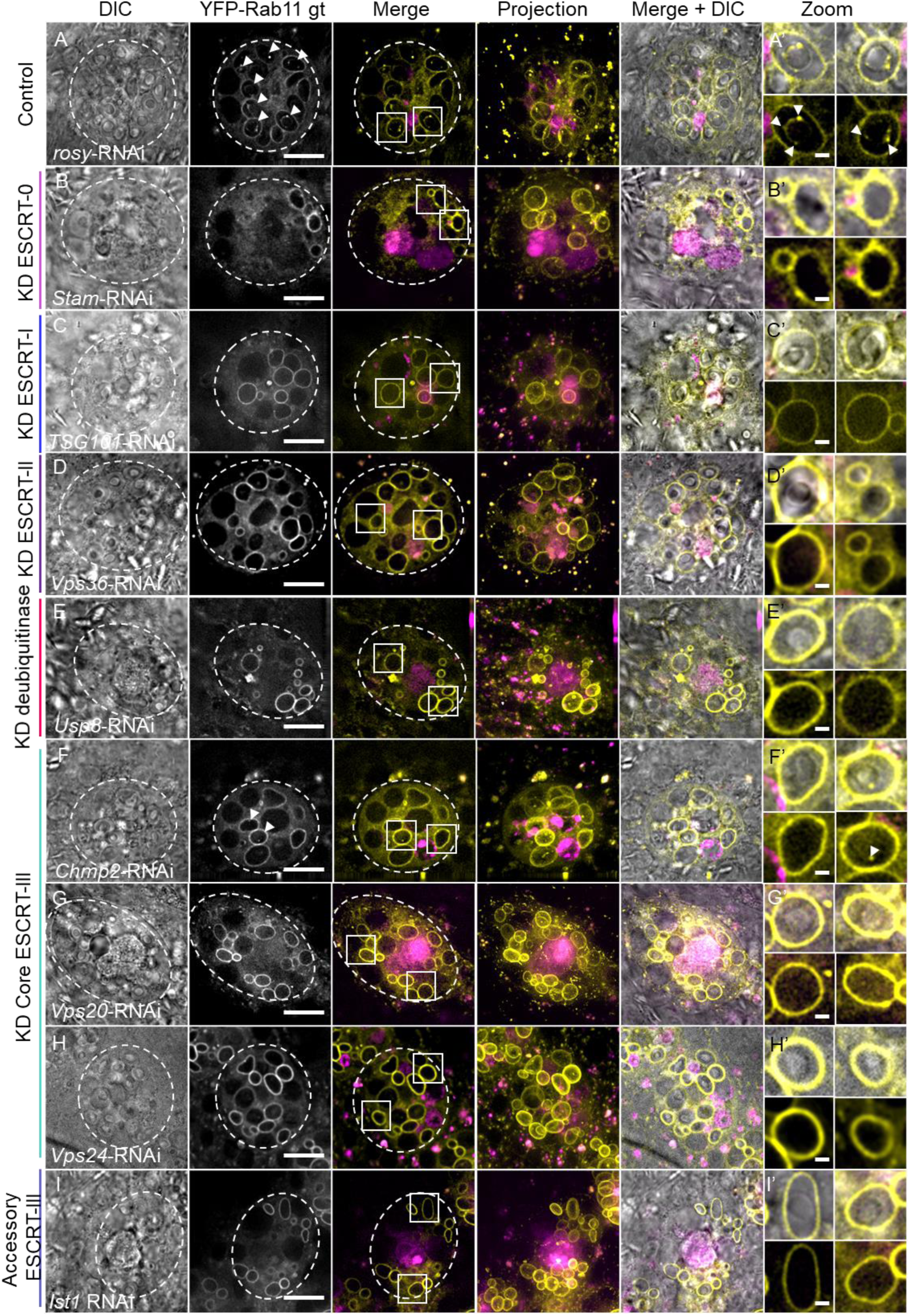
*ESCRT* knockdown inhibits exosome biogenesis in non-acidic SC compartments, but does not change their identity. Panels A-I show basal wide-field fluorescence views of living SCs expressing YFP-Rab11 from its endogenous genomic location (yellow). SC outline approximated by dashed white circles. Acidic compartments are marked by LysoTracker® Red (magenta, Merge). Boxed non-acidic compartments are magnified in A’-H’. (A) Control SC expressing *rosy-*RNAi. YFP-Rab11-positive ILV puncta are observed inside nearly 50% of Rab11-compartments (arrowheads; A’). (B) SC expressing *Stam-*RNAi. YFP-Rab11-positive ILVs are reduced. (C) SC expressing *TSG101-*RNAi. YFP-positive ILVs are reduced. (D) SC expressing *Vps36-*RNAi. YFP-Rab11-positive ILVs are reduced. (E) SC expressing *Usp8-*RNAi. YFP-Rab11-positive ILVs are reduced. (F) SC expressing *Chmp2-*RNAi. YFP-Rab11-positive ILVs are reduced. (G) SC expressing *Vps20-*RNAi. YFP-Rab11-positive ILVs are reduced. (H) SC expressing *Vps24*-RNAi. YFP-Rab11-positive ILVs are reduced. (I) SC expressing Ist1-RNAi. YFP-Rab11-positive ILVs are reduced. All data are from 6-day-old males shifted to 29°C at eclosion to induce transgene expression. Genotypes are: *w; P[w*^*+*^, *tub-GAL80ts]/+; dsx-GAL4/TI{Tl}Rab11EYFP* with RNAi #1 lines for each gene. Scale bars in A-I, 10 µm and in A’-I’, 1 µm.

**Supplementary Figure S4.**
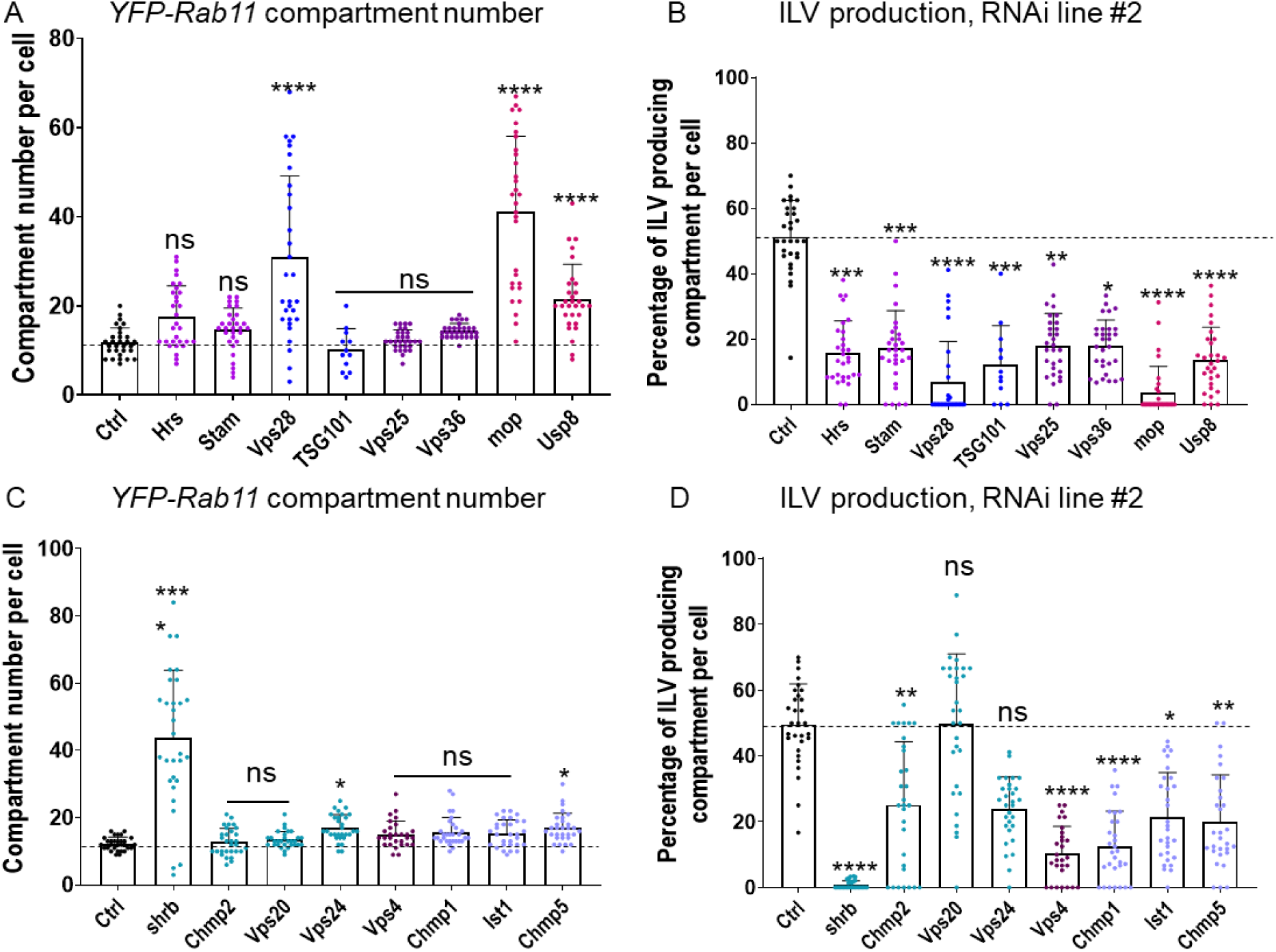
*ESCRT* knockdown inhibits exosome biogenesis in non-acidic SC compartments, but does not change their identity. Quantification of live SCs expressing RNAi #2 lines for all genes tested in Figs 4, 5 and S3. (A) Bar chart showing the number of non-acidic YFP-Rab11-positive compartments with diameter > 0.4 µm per SC in control vs ESCRT-0, -I, -II and deubiquitinases *mop* and *Usp8* knockdowns. n = 30. (B) Bar chart showing percentage of YFP-Rab11 compartments containing YFP-Rab11-positive ILVs. n = 30. (C) Bar chart showing the number of non-acidic YFP-Rab11-positive compartments with diameter > 0.4 µm per SC in control vs *ESCRT-III* and *Vps4* knockdowns. n = 30. (D) Bar chart showing percentage of YFP-Rab11 compartments containing YFP-Rab11-positive ILVs. n = 30. All data are from males shifted to 29°C at eclosion to induce transgene expression. Genotypes are: *w; P[w*^*+*^, *tub-GAL80ts]/+; dsx-GAL4/TI{Tl}Rab11EYFP* with *UAS-rosy-RNAi*, and RNAi #2 lines for each gene. Data were analysed by Kruskal-Wallis test. *p < 0.05, **p < 0.01, ***p < 0.001, and ****p < 0.0001 relative to control.

**Supplementary Figure S5.**
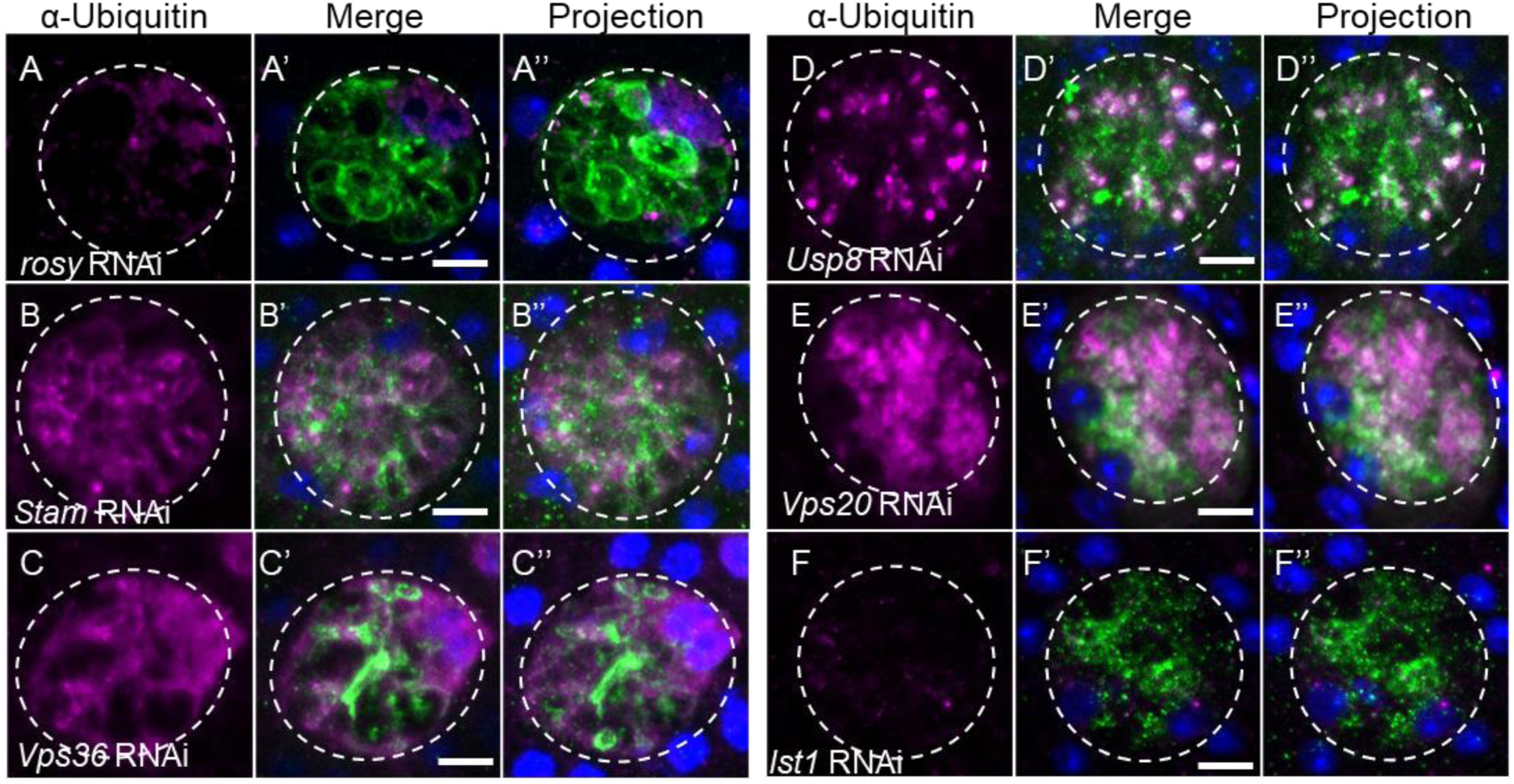
Accessory ESCRT-III proteins are not required for processing of ubiquitinylated ILV cargos. Additional confocal basal images of fixed SCs isolated from males expressing Btl-GFP and selected ESCRT-RNAis from eclosion onwards. SC outline approximated by dashed white circles. Ubiquitin (magenta), GFP (green) and DAPI (blue; marking binucleate cells) staining is shown. Nuclear staining with anti-ubiquitin is sometimes observed, even in controls, but appears non-specific. (A) Control SC expressing *rosy-* RNAi. Virtually no cytoplasmic accumulation of ubiquitin is observed. (B) SC expressing *Stam*-RNAi accumulates ubiquitin in cytoplasm. (C) SC expressing *Vps36*-RNAi shows some cytoplasmic accumulation of ubiquitin. (D) SC expressing *Usp8-* RNAi strongly accumulates ubiquitin in cytosol and some Btl-GFP-positive compartments. (E) SC expressing *Vps20-*RNAi accumulates ubiquitin in cytoplasm. (F) SC expressing *Ist1-*RNAi does not accumulate ubiquitin in the cytoplasm. All data are from 6-day-old male flies shifted to 29°C at eclosion to induce transgene expression. Genotypes are: *w; P[w*^*+*^, *tub-GAL80ts]/+; dsx-GAL4/P[w*^*+*^, *UAS-btl-GFP]* with *rosy-* or RNAi #1 lines. Scale bars in A-F, 10 µm.

